# Deep homology of a *brachyury* regulatory syntax and origin of the notochord

**DOI:** 10.1101/2024.12.12.628285

**Authors:** Tzu-Pei Fan, Jun-Ru Lee, Che-Yi Lin, Yi-Chih Chen, Ann E. Cutting, R. Andrew Cameron, Jr-Kai Yu, Yi-Hsien Su

**Author notes:** Correspondence: Yi-Hsien Su. Deceased November 19, 2022 (R. Andrew Cameron).

## Abstract

The *brachyury* gene encodes a T-box transcription factor (TF) that is crucial for the development of the notochord, a novel trait of chordates^1^. *Brachyury* expression in axial mesodermal cells (notochord progenitors) is regarded as a chordate innovation^2^, yet it remains unclear how the chordate ancestor acquired this expression domain. By examining the sequences of previously identified notochord enhancers of several chordate *brachyury* genes^3–5^, we uncovered a regulatory syntax consisting of binding sites for four TFs (Su(H), Foxh1, Zic, and Ets) with a strict order and orientation. We also identified this syntax, here named SFZE, in potential *cis*-regulatory modules (CRMs) of *brachyury* orthologs in various non-chordate animals and even in *Capsaspora*, a close unicellular relative to animals. Reporter assays demonstrated that SFZE-containing CRMs from non-chordate organisms exhibited regulatory activity in the zebrafish notochord, and mutagenesis of the TF sites reduced the activity. Furthermore, the SFZE syntax in sea urchin confers its endoderm activity, with the TF sites functionally decoupled during gastrulation. These findings indicate that the association of the SFZE syntax with *brachyury* is ancient, likely predating the origin of animals. The emergence of axial *brachyury* expression is therefore probably not the result of a newly acquired notochord enhancer, but is instead likely attributed to co-option of upstream signals acting on the conserved SFZE syntax, which facilitates the origin of the notochord from rudimentary endodermal cells.

---

The notochord is a defining feature of chordates (including amphioxus, tunicates, and vertebrates) that plays critical roles in chordate development. The notochord cells are differentiated from the axial mesoderm, which arises from the dorsal organizer located on the dorsal side of the blastopore at the onset of gastrulation. As gastrulation proceeds, the dorsal organizer and its derived notochord send signals to pattern the surrounding tissues and construct the basic chordate body plan^6^. The two closest relatives of chordates are the hemichordates and echinoderms (together called ambulacrarians); these three groups constitute the deuterostomes. It has been proposed that the hemichordate stomochord, an anterior rod-like protrusion from the pharynx, is homologous to the chordate notochord. However, molecular studies do not support stomochord-notochord homology^7^. In a protostome annelid, it was shown that a population of midline mesodermal cells expresses a set of notochord-specific genes and differentiates into a medial ventral longitudinal muscle named the axochord^8^. While the axochord has been proposed to be a notochord homolog, the annelid axochord is formed days after gastrulation, and it remains possible that the two structures may have evolved convergently^9^.

*Brachyury* encodes a T-box TF that is crucial for notochord development^10,11^. The *brachyury* gene is considered to be the most ancient among the T-box family, and its orthologs have been identified in various metazoans and also in several non-metazoan lineages (such as protists and fungi)^12,13^. *Brachyury* genes from non-chordate metazoans transform ascidian endodermal cells into notochord cells^14^, suggesting that Brachyury proteins of non-chordates and chordates are functionally equivalent. In contrast to the conservation of protein function, the *brachyury* expression patterns in chordates differ from those of non-chordates. In chordates, the expression pattern of *brachyury* is generally conserved. For example, *brachyury* transcripts in amphioxus and frog are first detected in a ring of cells surrounding the blastopore. Subsequently, the *brachyury*-expressing cells internalize during gastrulation and differentiate into the notochord^15,16^. In various non-chordate bilaterians, a circumblastoporal expression domain is also observed in gastrulae; *brachyury* is also expressed in the hindgut and oral ectoderm, but never in axial mesodermal cells^17^. These observations have led to a hypothesis that the circumblastoporal expression of *brachyury* is an ancestral trait in bilaterians, while *brachyury* expression in the axial mesoderm represents a novelty that arose in the chordate ancestor^2^. Therefore, the evolutionary origin of the notochord is likely associated with the gain of a novel *brachyury* expression domain in the axial mesoderm, but not with functional changes of the Brachyury protein^2^. Emergence of new expression domains is often attributed to births or modifications of CRMs. Previous studies in the ascidian *Ciona* and zebrafish have identified CRMs upstream of the *brachyury* translation start site (TSS) that are able to drive reporter gene expression in the notochord^4,5,18^. It remains to be determined whether these notochord enhancers evolved *de novo* in the chordate ancestor or if they have a more ancient origin.

## Ambulacrarian *brachyury* BACs are active in zebrafish

To explore the evolutionary origin of the *brachyury* notochord enhancer, we aimed to dissect CRMs of *brachyury* genes from non-chordate animals and test their activities in chordates. We first focused on animals that are closely related to chordates, including the hemichordate *Ptychodera flava* and the sea urchin *Strongylocentrotus purpuratus*. We generated *gfp* knock-in BACs (bacterial artificial chromosomes) harboring the *brachyury* ortholog of either *P. flava* or *S. purpuratus* (*Pfbra*/*Spbra*:*gfp* BACs). Injections of the reporter BACs into their cognate embryos recapitulated the endogenous expression of *brachyury* in the blastopore, oral ectoderm, and endoderm^19,20^ (Extended Data Fig. 1 and 2), although some ectopic expression was also observed. These results suggest that the BACs contain CRMs sufficient to drive *brachyury* expression at the analyzed stages. To examine CRM activities of the ambulacrarian *brachyury* genes in chordate embryos, we introduced each of the reporter BACs into zebrafish zygotes (Fig. 1a). At the onset of gastrulation (shield stage, 6.5 hpf), approximately 33% of the zebrafish embryos injected with the *Pfbra*:*gfp* BAC showed GFP signals in the embryonic shield (Fig. 1b, Extended Data Table 2). This structure is equivalent to the dorsal organizer that gives rise to midline structures including the notochord^21,22^. At the early segmentation stage (11-14 hpf), GFP signal was detected in the dorsal midline in ∼45% of the transgenic embryos (Fig. 1c-1d, Extended Data Table 2). In contrast to the strong reporter activity driven by *Pfbra*:*gfp* BAC, GFP signal was weak or not observed in most of the *Spbra*:*gfp* BAC-injected zebrafish embryos. Out of the 174 injected embryos, only two displayed GFP signals in the organizer at the shield stage (Fig. 1e, Extended Data Table 2), and the signals became faint at the early segmentation stage (Fig. 1f-1g). The detected GFP domains in the shield and the dorsal midline before and after gastrulation resemble the developmental progression of the notochord, which can be tracked by the expression of *Drntl* (*no tail*), a zebrafish ortholog of *brachyury*^23^. To determine whether *gfp* was indeed activated in the *Drntl*-expressing cells, we performed double fluorescent *in situ* hybridization (dFISH). At the shield stage, *Drntl* transcripts were detected in a ring at the margin (germ ring), and *gfp* transcripts were observed in the *Drntl*-expressing cells in the shield (Fig. 1h-1k). At the early segmentation stage, *gfp* transcripts were detected in the dorsal midline underneath the *Drntl* expression domain, but not in the *Drntl*-expressing notochord cells (Fig. 1l-1p).

**Figure 1.**
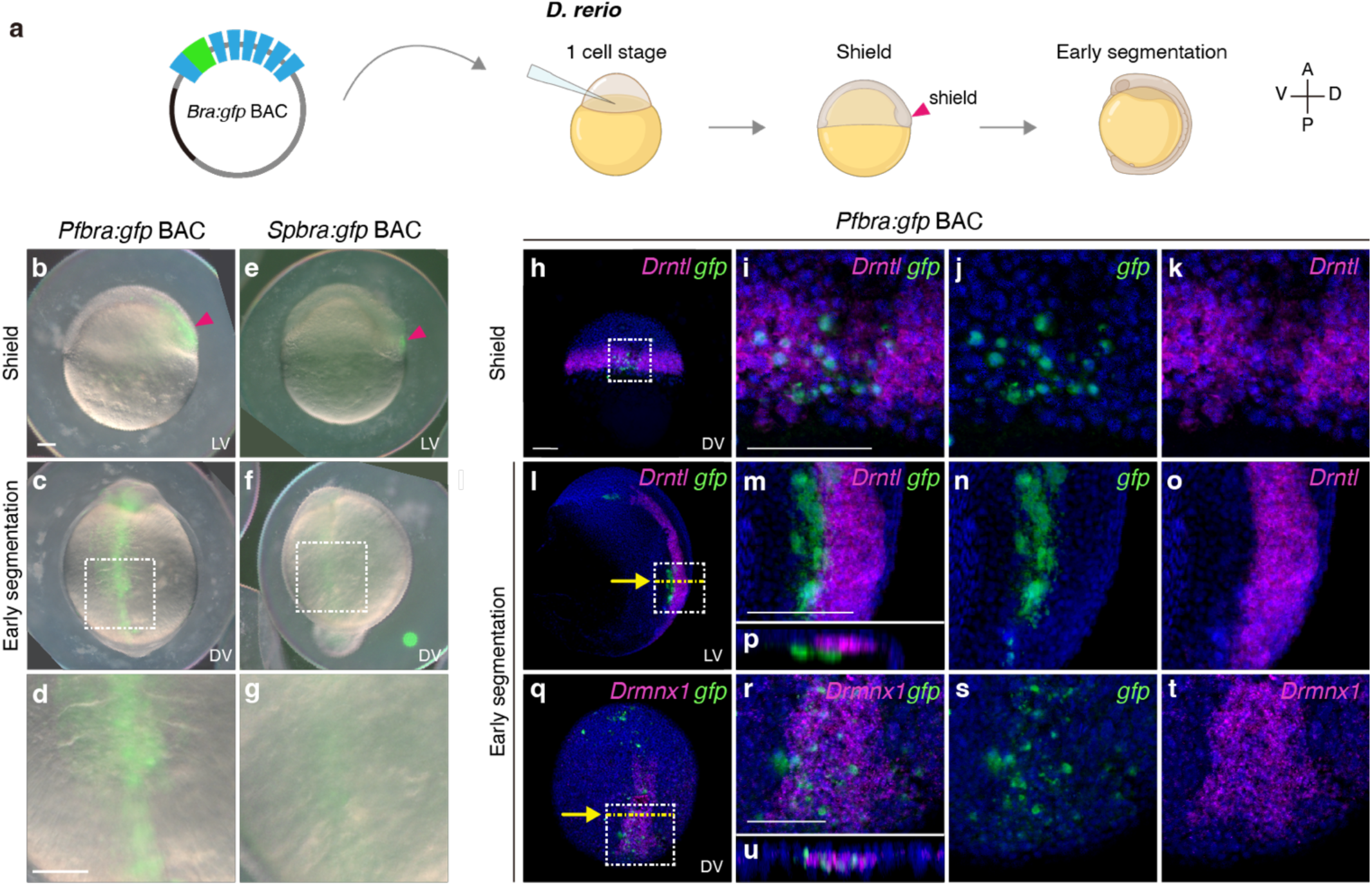
*Pfbra:gfp* BAC activates GFP expression in the organizer and the hypochord of zebrafish embryos. **a,** Circular *Pfbra*/*Spbra*:*gfp* BACs were each introduced into zebrafish zygotes, and GFP signals were observed at the shield and early segmentation stages (A, anterior; P, posterior; V, ventral; D, dorsal). **b-g,** GFP signals in zebrafish embryos injected with *Pfbra*:*gfp* BAC or *Spbra*:*gfp* BAC. GFP signal in the dorsal margin (shield) at the shield stage is indicated by the red arrowhead. The white dashed boxes in panels c and f are magnified and shown in panels d and g, respectively. **h-u,** Double FISH of *gfp* (green) and *Drntl* (magenta in panels h-p) or *Drmnx1* (magenta in panels q-u) in *Pfbra:gfp* BAC-injected zebrafish embryos at indicated stages. Nuclei were counterstained with Hoechst 33342 (blue). The areas in the white dashed boxes in panels h, l, and q are enlarged and shown in panels i, m, and r; single-channel images are shown in panels j-k, n-o, and s-t, respectively. **p and u,** The XZ sections along the yellow dashed lines are indicated with yellow arrows in panels l and q. The orientation of embryos is indicated in each panel: LV, lateral view; DV, dorsal view. All scale bars represent 100 μm. Panels that are in the same scale: b-c and e-f; d and g; h, l, and q; i-k, m-p, and r-u. Illustrations of zebrafish embryos were created based on images from BioRender.com.

The GFP signal in cells ventral to the notochord likely marked the hypochord, a midline structure sharing the same progenitors with the notochord^22,24^. This localization was confirmed by dFISH analysis, which showed *gfp* was expressed in hypochord cells marked by the expression of *Drmnx1*^25^ (Fig. 1q-1u and Extended Data Fig. 3). Together, these results indicate that the ambulacrarian *brachyury* CRMs respond to the regulatory state of the notochord progenitors in the shield, but not in the notochord at the later segmentation stage. Previous studies have shown that the *Drntl*-expressing cells in the organizer are bipotential for notochord and hypochord^22,24^, and their fate decisions depend on Delta signal from the adjacent cells.^22^. Our data thus suggest that, while *Pfbra*:*gfp* BAC is transcriptionally active in the zebrafish notochord progenitors, it may lack an enhancer to maintain its activity or possess a silencer(s) to suppress reporter expression in the developing notochord.

## Hemichordate PfCRM2 is active in the notochord

To identify CRM(s) of *Pfbra* that contribute to the transcriptional activities in zebrafish embryos, we analyzed published ATAC-seq data^26^ seeking to identify open chromatin regions at the *Pfbra* locus (Fig. 2). Among the four major ATAC-seq peaks representing potential CRMs, PfCRM2 spans 717 bp immediately upstream of the TSS and contains the promoter (Fig. 2s). We generated reporter constructs of PfCRM2 and the other three PfCRMs fused with the deduced *Pfbra* promoter (a 332 bp fragment upstream of the TSS), using an *egfp* vector^27^ (Extended Data Fig. 4a). Reporter constructs with *Pfbra* promoter alone or containing the known notochord enhancer of *Drntl* (∼1 kb upstream of the *Drntl* TSS, *Drntl*-1kb)^4^ were also generated for comparison (Extended Data Fig. 4a). At the shield stage, *Drntl*-1kb and PfCRM2 drove *egfp* expression in the hypoblast of the embryonic shield, the precursor of the notochord, in the majority of the embryos (on average, 98% for *Drntl*-1kb and 80% for PfCRM2). EGFP signals were also observed in the dorsal germ ring for some embryos (Extended Data Fig. 4b-4c and 4h-4i, Extended Data Table 3). Reporters with the *Pfbra* promoter alone or with PfCRM1, PfCRM3, or PfCRM4 exhibited lower activities in the hypoblast (Extended Data Fig. 4d-4n, Extended Data Table 3). At the early segmentation stage, *Drntl*-1kb and PfCRM2 constructs showed notochord activity in 49% and 33% of EGFP-positive embryos, respectively (Fig. 2b-2e, 2x). In contrast, we observed very few or no embryos with EGFP signal in the notochord after injection with reporters containing *Pfbra* promoter alone or with PfCRM1, PfCRM3 or PfCRM4 (Fig. 2x, Extended Data Fig. 4o-4v, Extended Data Table 3). We further performed dFISH to confirm that the EGFP-positive cells in the PfCRM2-injected embryos were indeed *Drntl*-expressing notochord cells (Extended Data Fig. 4w-4z). These results indicate that among the four PfCRMs, only PfCRM2 exhibits comparable transcriptional activities in the hypoblast and notochord to those of *Drntl*-1kb. Thus, it appears that PfCRM2 shares similar regulatory features with the zebrafish notochord enhancer. Additionally, the *Pfbra* promoter possesses limited regulatory function in the hypoblast, and it likely accounts for the hypoblast activities of PfCRM1 and PfCRM4. Furthermore, the observation that PfCRM2, but not *Pfbra*:*gfp* BAC, is transcriptionally active in the notochord implies the presence of silencer(s) within the BAC that suppress PfCRM2 activity in the notochord. PfCRM3 likely serves as a silencer, given that it significantly reduced *Pfbra* promoter activity in the hypoblast and the absence of its activity in the notochord.

**Figure 2.**
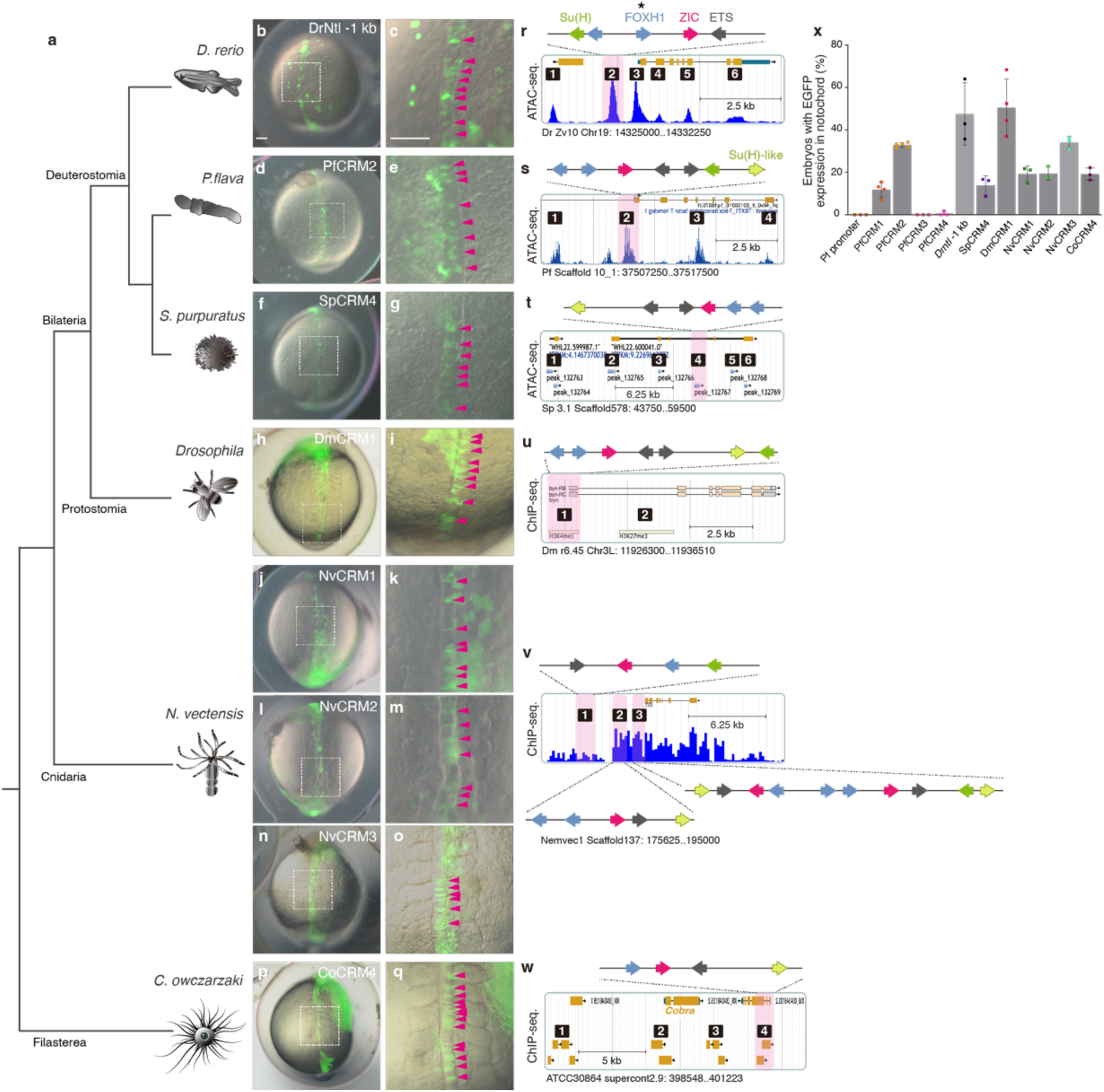
*Brachyury* CRMs with SFZE/SFZE-like syntax show activity in the zebrafish notochord. **a.** Phylogenetic relationships of the analyzed species. **b-q**, Zebrafish embryos injected with SFZE/SFZE-like containing CRMs from zebrafish (*Drntl*-1 kb), hemichordate (PfCRM2), sea urchin (SpCRM4), fruit fly (DmCRM1), sea anemone (NvCRM1, NvCRM2 and NvCRM3), and filasterea (CoCRM4). The right panels show magnified views of the notochord region (white dashed boxes in the corresponding left panels) with magenta arrowheads indicating EGFP signals in the notochord cells. The left panels are in the same scale, as are the right panels. Scale bars in panels b and c, 100 μm. **r-w**, The putative CRMs based on published ATAC-seq (*Drntl*, *Pfbra*, and *Spbra* loci) and ChIP-seq datasets (*Dmbyn*, H3K4me1 and H3K27me3; *Nvbra*, H3K4me1; *Cobra*, H3K4me1 and H3K4me3). Potential CRMs are indicated with numbers, and CRMs carrying SFZE/SFZE-like motifs are shaded in pink. The TF binding sites are denoted schematically with colored arrows placed on top of each diagram to show the orientation and order. Colors of TF sites are shown in panels r and s. Genome versions, scaffold/chromosome numbers, and positions of the genomic loci displayed in panels r-w are indicated below the respective panel. The samples used for generating of the ATAC-seq and ChIP-seq datasets are listed in Extended Data Table 7. **x**, Percentages of embryos with EGFP signals in the notochord out of EGFP-positive embryos. Each colored square represents the result of one experiment. Gray columns represent the average of at least three biological replicates, with error bars serving as standard deviations.

## SFZE syntax of *brachyury* notochord enhancers

To identify the regulatory features that may account for the transcriptional activity in the notochord, we compared sequences of PfCRM2 with the known notochord enhancers of *brachyury* orthologs from two chordate species, zebrafish^4^ and the ascidian *Ciona intestinalis type A* (or renamed as *Ciona robusta*)^18,28^. Using mVISTA^29^, we were unable to detect conserved regions between PfCRM2 and the notochord enhancers (Extended Data Fig. 5). Nevertheless, the ability of PfCRM2 to drive reporter gene expression in the notochord prompt us to hypothesize that TF binding sites within PfCRM2 are sufficient to respond to particular TFs present in the zebrafish notochord. Studies to dissect the regulatory controls of zebrafish and *Ciona brachyury* genes have identified TF binding sites that are critical for expression in the notochord. These include a Foxh1 (also known as Fast-1) binding site located within *Drntl*-1kb^4^ and two binding sites for Suppressor of Hairless (Su(H)) within the proximal enhancer of the *Ciona brachyury* gene (*Cibra*)^18^; one complies the known consensus sequence (GTGRGAR)^30^, while the other is less ideal (NTGRGAR). Additionally, a shadow enhancer located ∼800 bp upstream of the *Cibra* TSS has been found to drive reporter gene expression in the notochord. This enhancer contains two Ets sites and one ZicL site; of note, the face-to-face orientation of the Ets and ZicL binding sites is critical for enhancer function^3^. A study in another ascidian, *Halocynthia roretzi*, showed that both sites of Ets and ZicN in the promoter region of the *brachyury* gene (*Hrbra*) are required for *Hrbra* initiation in notochord precursors^5^. Based on these published results, we reasoned that rather than sequence conservation, TF sites with specific grammar may be essential for transcriptional activity in the notochord. The known notochord enhancer of *Drntl* encompasses two ATAC-seq peaks^31^ (DrCRM2 and DrCRM3 in Fig. 2r). Within DrCRM2, we found binding sites for Zic, Ets and Su(H) adjacent to the previously identified Foxh1 site^4^ (Fig. 2r and Extended Data Fig. 6); an additional Foxh1 site was also observed. Importantly, similar to the orientation found in the shadow enhancer of *Cibra*^3^, the Zic and Ets sites of DrCRM2 are facing each other (Fig. 2r, Extended Data Fig. 6). We further examined sequences of the *Cibra* shadow enhancer and uncovered two Foxh1 binding sites adjacent to the previously identified ZicL site (Extended Data Fig. 6, Extended Data Fig. 7). In sum, the notochord enhancers of both zebrafish and *Ciona brachyury* genes contain binding sites of the four TFs distributed in a specific order (5’-Su(H)-Foxh1-Foxh1-Zic-Ets-3’ for zebrafish and a reverse order for *Ciona*) and with specific orientations (face-to-face orientation of Zic and Ets sites, and the same direction for Zic and the adjacent Foxh1 site). These TF sites are also in close proximity to each other, especially the Foxh1-Zic-Ets sites (span less than one nucleosome length, ∼147 bp; Extended Data Table 4). These features suggest that the sites may function collaboratively in recruiting TFs to regulate *brachyury* expression^32^. Hereafter, we use ‘SFZE’ (based on the first letter for each of the four TFs) to refer to the syntax of the binding sites, including the specific order, orientations and spacing. We postulated that the SFZE syntax likely represents a functional unit for driving *brachyury* expression in the notochord. In line with this hypothesis, we were unable to identify the SFZE syntax in the *Xenopus brachyury* (*Xbra*) promoter, which activates reporter gene expression in the blastopore and not in the notochord^33^. Together, our results to this point suggested that the SFZE syntax is likely a conserved feature in the notochord enhancer of chordate *brachyury* genes.

## Deep homology of the SFZE syntax

To explore whether the SFZE syntax has an ancient origin and might account for the notochord activity of the hemichordate PfCRM2, we scanned PfCRM sequences for binding sites of the four TFs. We discovered that PfCRM2 contains two Foxh1 sites, followed by one Zic, two Ets, and two Su(H) sites (one is a less optimal Su(H)-like site; Fig. 2s, Extended Data Fig. 6). These sites are closely associated and comply with the chordate SFZE syntax, as the Zic and Ets sites face each other, and the Foxh1 site is oriented in the same direction as the adjacent Zic site. One difference is that the position of one Su(H) site is adjacent to Ets sites in the hemichordate but to Foxh1 sites in the zebrafish SFZE syntax. Thus, the notochord activity of PfCRM2 is likely attributed to the presence of the SFZE syntax, with FZE serving as the core of the syntax and the position of Su(H) sites being relatively less critical. Importantly, PfCRM3 does not contain Ets and Su(H) sites, and PfCRM4 lacks Su(H) sites, consistent with the absence of notochord activity (Fig. 2x, Extended Data Fig. 6). PfCRM1 contains binding sites for all four TFs, but the Ets and Zic sites are oriented in the same direction (Extended Data Fig. 6). Consistent with this slightly disrupted syntax, PfCRM1 exhibited lower notochord activity than PfCRM2, but higher activity than PfCRM3 and PfCRM4 (Fig. 2x). We further examined the sequences of potential CRMs of the sea urchin *Spbra* gene. Among the putative CRMs uncovered in available ATAC-seq data^34^, we identified an SFZE motif in SpCRM4, which is located in the third intron of the gene (Fig. 2t). The sea urchin SFZE motif complies with the FZE orientation rule, but is partially inverted compared to the SFZE syntax (i.e., SEZF, Extended Data Fig. 6). Nevertheless, SpCRM4 exhibited transcriptional activity in the zebrafish notochord, albeit at a lower level (Fig. 2f-g, 2x, Extended Data Table 5). Together, these results strongly suggest the role of the SFZE syntax in the notochord activity and demonstrate that the association of the SFZE syntax with *brachyury* orthologs is present in non-chordate deuterostomes, predating the origins of the notochord.

To further trace the origin of the SFZE syntax, we analyzed potential CRMs associated with the *brachyury* ortholog (*brachyenteron*) of *Drosophila melanogaster* (*Dmbyn*), a representative species of protostomes (Fig. 2a). Using an available ChIP-seq dataset^35^ that shows two putative enhancer regions, we identified the SFZE syntax in DmCRM1 reporter at the *Dmbyn* locus (Fig. 2u). The order of TF binding sites in DmCRM1 is exactly the same as that in the hemichordate PfCRM2 (Fig. 2u, Extended Data Fig. 6). Furthermore, the DmCRM1 was able to drive *egfp* expression in the zebrafish notochord (Fig. 2h-2i), and the percentage of embryos exhibiting EGFP signals in the notochord was comparable to that of embryos injected with the zebrafish reporter (Fig. 2x and Extended Data Table 5). These results reinforce the idea that the notochord activity is attributed to the SFZE syntax. Moreover, the results suggest that the SFZE syntax has a deep evolutionary history, likely originating before the divergence of protostomes and deuterostomes, and representing a conserved feature of bilaterian *brachyury* genes.

To examine whether non-bilaterian animals possess the SFZE syntax, we analyzed the sequences of three potential CRMs surrounding the *brachyury* ortholog of the sea anemone *Nematostella vectensis* (*Nvbra*) (Fig. 2v). Analysis of the available ChIP-seq dataset^36^ revealed that NvCRM1 contains an inverted FZE motif upstream of a Su(H) site. Within NvCRM2, the Ets and Zic sites are oriented in the same direction, while the Zic and the adjacent Foxh1 sites are not in the same direction, violating the FZE orientation rules. NvCRM3 has two consecutive SFZE motifs. The upstream motif has an inverted FZE syntax, identical to the motif order within the sea urchin SpCRM4. Meanwhile, the downstream motif does not conform to the FZE grammar rules, with Ets and Zic sites oriented in the same direction. Furthermore, the Ets, Zic, and Fox sites within the upstream SFZE motif in NvCRM3 are positioned in close proximity to each other (within one nucleosome length; Extended Data Table 4), similar to what we observed in bilaterians. In contrast, these sites in NvCRM1, NvCRM2, and the downstream SFZE motif of NvCRM3 span more than one nucleosome length (Extended Data Table 4). Consistent with the presence of the SFZE syntax and the close association of the FZE sites, the NvCRM3 reporter exhibited higher transcriptional activity in the zebrafish notochord than the NvCRM1 and NvCRM2 constructs (Fig. 2j-2o, 2x, and Extended Data Table 5). In fact, NvCRM3 activity was even higher than that of the sea urchin SpCRM4, likely due to a synergistic effect of the two SFZE motifs. The deeply conserved SFZE syntax within animals led us to further trace its association with the *brachyury* ortholog of the filasterean *Capsaspora owczarzaki* (*Cobra*)^12^, a unicellular eukaryote that is phylogenetically close to metazoans. Within one potential CRM (CoCRM4) identified from a published ChIP-seq dataset^37^, we found an SFZE syntax containing one Su(H)-like site with the FZE core located within an exon of a gene (CAOG_05510) neighboring *Cobra* (CAOG_05512) (Fig. 2w; Extended Data Fig. 6). Importantly, this non-animal CoCRM4 was also able to drive *egfp* expression in the zebrafish notochord (Fig. 2x and Extended Data Table 5). Together, these results showed the association of the SFZE syntax with *brachyury* orthologs of two ambulacrarians (hemichordate and sea urchin), a protostome (*Drosophila*), a cnidarian (*Nematostella*), and a unicellular relative of animals (*Capsaspora*). Furthermore, reporter constructs containing CRMs with SFZE syntax exhibited transcriptional activities in the zebrafish notochord. Our results thus strongly support the conclusion that the SFZE syntax is deeply conserved, and it likely originated before the emergence of animals.

## Foxh1 and Ets sites are functionally important

Activation of zebrafish *brachyury* in the notochord is known to rely on Nodal and FGF signals, which respectively act via Foxh1 and Ets TFs^4,38–40^. To evaluate whether the Foxh1 and Ets sites within the hemichordate SFZE syntax are also critical, we removed the predicted binding sites in PfCRM2. When the Foxh1 sites were deleted, the reporter activities in the zebrafish notochord were significantly reduced, but not completely abolished (Fig. 3a-3e, 3h). In contrast, mutation of the two Ets sites strongly decreased reporter activity in the notochord (Fig. 3f-3g, 3h). These results demonstrate that both Foxh1 and Ets sites in the SFZE of PfCRM2 are functional, suggesting that these sites may be capable of responding to the endogenous Nodal and FGF signaling pathways in the notochord. Additionally, the Ets sites in the SFZE motif play a crucial role, while the Foxh1 sites seem to confer an additive effect, strengthening the transcriptional activity in the notochord.

**Figure 3.**
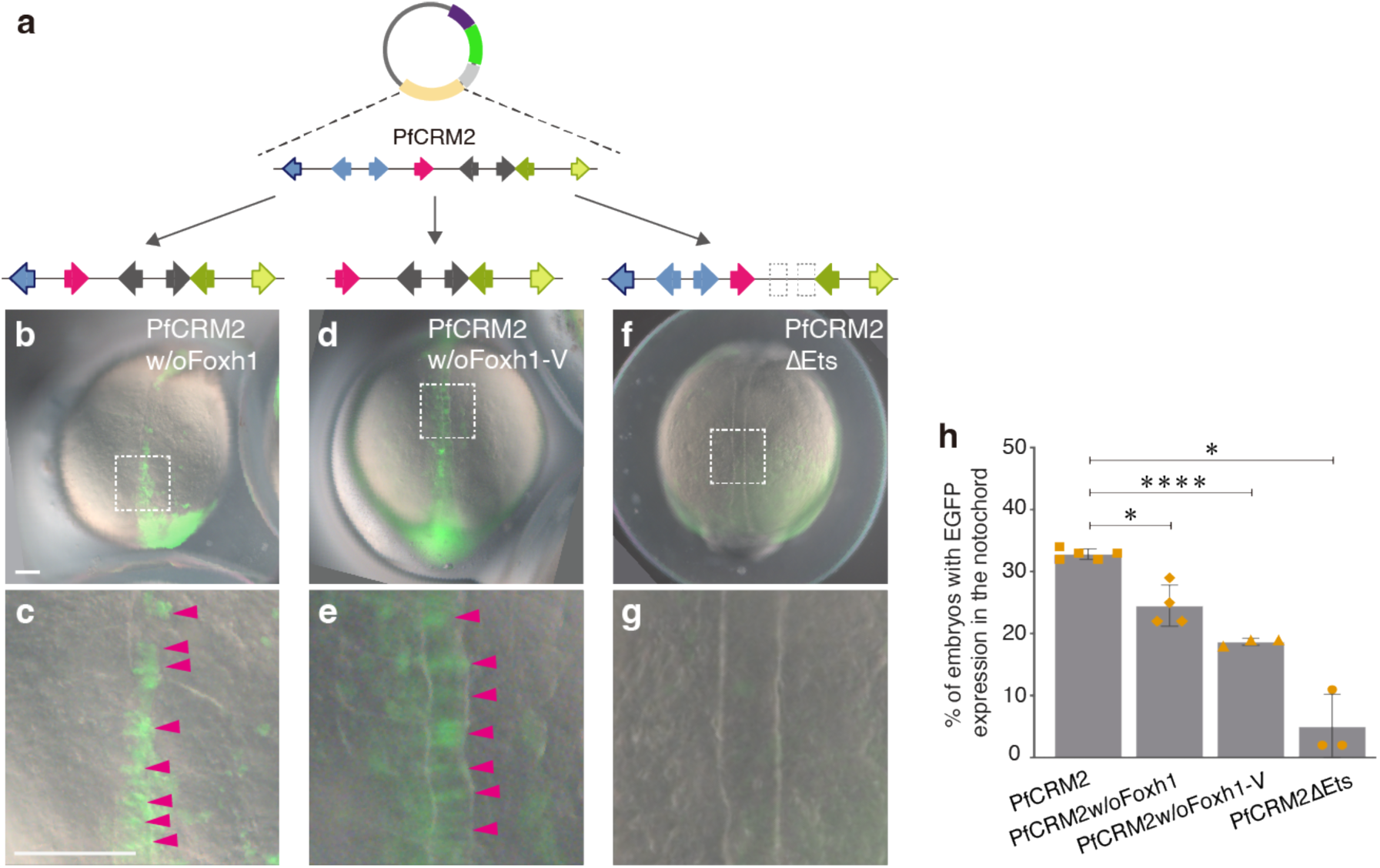
Foxh1 and Ets sites of the hemichordate SFZE syntax are functionally important for notochord activity. **a**, Diagram of the reporter constructs of PfCRM2 with deletions of the Foxh1 sites or mutations of the Ets sites. The additional Foxh1 site in the vector is indicated by the blue arrow with black outline. **b-g**, Representative images show zebrafish embryos injected with the truncated or mutated PfCRM2 reporter constructs. Magnifications of the notochord regions highlighted by white boxes in panels b, d and f are shown in panels c, e and g, respectively. EGFP signals in the notochord cells are indicated with magenta arrowheads. Embryos are oriented in the dorsal view with anterior to the top. Upper panels (b, d, and f) and lower panels (c, e, and g) are in the same scale as indicated in panels b and c, respectively. All scale bars represent 100 μm. **h**, Percentages of zebrafish embryos exhibiting EGFP signals in the notochord among EGFP-positive embryos. Each yellow data point represents the result of a single experiment. The gray columns are average results from at least three biological replicates, with error bars showing standard deviations. *: *p* < 0.05; ****: *p* < 0.0001.

## Sea urchin SFZE syntax confers endoderm activity

To this point, we had shown that non-chordate *brachyury* CRMs containing the SFZE syntax possess notochord activity in zebrafish embryos; however, their endogenous activities remained unknown. To explore the endogenous function of the SFZE syntax in organisms lacking a notochord, we analyzed the activity of SpCRM4 during sea urchin gastrulation (Fig. 4a). The *Spbra* promoter alone showed negligible background activity at the mesenchyme blastula (i.e. initiation of gastrulation) and late gastrula stages (Extended Data Table 6). The promoter driven by SpCRM4 exhibited strong activity in the presumptive endoderm at the mesenchyme blastula stage (Fig. 4b, 4j), recapitulating the endogenous expression pattern of sea urchin *brachyury* (Extended Data Fig. 8a-d). At the late gastrula stage, the SpCRM4 reporter was active in the archenteron, non-oral ectoderm, cells surrounding the blastopore, and mesenchymal cells (Fig. 4e and 4k). The ectodermal pattern is somewhat different from that of the *Spbra*:*gfp* BAC, which recapitulates the endogenous *brachyury* expression in the oral ectoderm (Extended Data Fig. 2, Fig. 4k, and Extended Data Fig. 8e-g). Additional CRMs within the *Spbra*:*gfp* BAC may account for this difference in ectodermal expression. To examine whether the Foxh1 and Ets sites of the SFZE syntax in SpCRM4 also contribute to its transcriptional activity, we generated reporter constructs with mutated Foxh1 or Ets sites. At the mesenchyme blastula stage, mutation of the two Foxh1 binding sites had no effect, while disruption of the two Ets sites decreased the activity in the presumptive endoderm moderately but significantly (Fig. 4c-d and 4j). At the late gastrula stage, mutation of the two Foxh1 sites decreased EGFP signals in the archenteron moderately but significantly, whereas mutated Ets sites had little effect (Fig. 4f-4i and 4k). These results led us to conclude that SpCRM4 functions mainly as an endoderm enhancer at the onset of gastrulation, and it behaves as a general enhancer in all three germ layers as gastrulation proceeds. Moreover, the endodermal activities of SpCRM4 at the onset and at the end of gastrulation partially depend on the respective Ets and Foxh1 sites within the SFZE syntax. Given that the endodermal expression patterns of *brachyury* orthologs are comparable in sea urchin, sea star and hemichordate (Extended Data Fig. 8), the endodermal activity of the SFZE syntax is likely a conserved feature in ambulacrarians.

**Figure 4.**
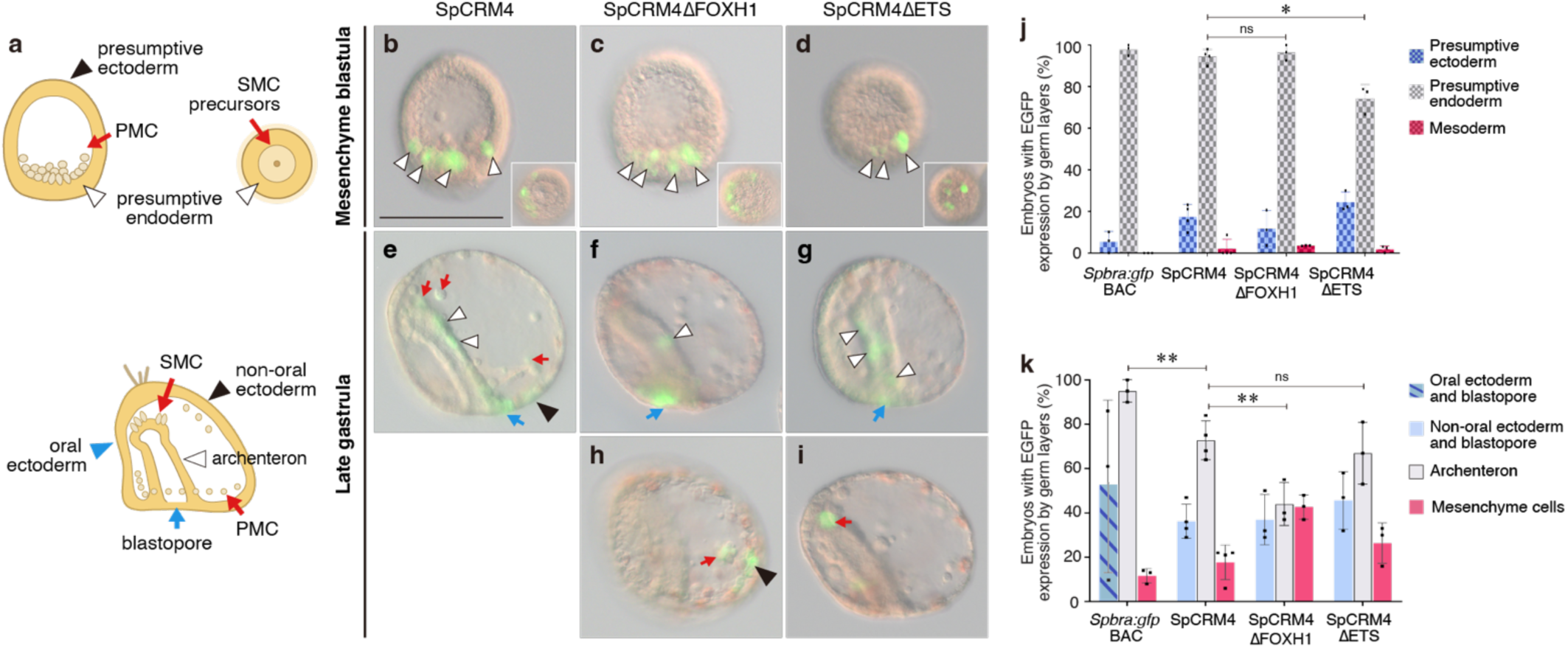
Foxh1 and Ets sites of the sea urchin SFZE syntax confer endodermal activity at different developmental stages. **a**, Illustrations of sea urchin embryos at the mesenchyme blastula (upper left: lateral view; upper right: vegetal view) and late gastrula (bottom) stages. Colored arrows and arrowheads mark different embryonic territories. **b-i**, Representative EGFP-positive embryos showing reporter activities. Colored arrows and arrowheads indicate EGFP-positive cells within the territories shown in panel a. Embryos are oriented as in panel a. For late gastrulae, the ventral side is to the left. Inserts in panels b-d are the vegetal view to show EGFP signals in the presumptive endoderm. Panels are all in the same scale (scale bar in panel b, 100 μm). **j-k**, Percentages of mesenchyme blastula (j) and late gastrula (k) embryos showing EGFP signals in the indicated territories. PMC, primary mesenchyme cells; SMC, secondary mesenchyme cells. *: *p* < 0.05; **: *p* < 0.01; ns: no significance.

## SFZE syntax in vertebrate *brachyury* paralogs

*Brachyury* is generally present as a single copy gene in invertebrates^41^. With two rounds of whole genome duplications (2R) during vertebrate evolution^42^ following lineage-specific loss of *brachyury* paralogs, ray-finned fishes have *tbxta* and *tbxtb* genes^43^, while tetrapods only retain *tbxtb*^41,43^. Given that the association of SFZE syntax with *brachyury* occurred before 2R, we anticipate that both vertebrate *brachyury* paralogs would have inherited SFZE motifs. The zebrafish SFZE we identified is associated with *tbxta* (i.e., *Drntl*). At the zebrafish *Drbra*/*tbxtb* locus, we uncovered the SFZE syntax in two putative CRMs situated 4.8 kb upstream and 12 kb downstream (Extended Data Fig. 6 and 7). Additionally, we found that a proximal CRM (1.6 kb upstream of the TSS) of the mouse *brachyury* (*Mmbra*/*T*/*tbxtb*) contains the SFZE syntax (Extended Data Fig. 6 and 7). Intriguingly, this CRM is located within the 8.3 kb region upstream of the TSS, which is able to direct *Mmbra* expression in the primitive streak (homologous to the blastopore) but not in the notochord of the mouse embryo^44^. We further analyzed the *Xenopus* genome using published ChIP-seq data^45^. At the *Xbra*/*tbxtb* locus, we found two putative CRMs containing the SFZE syntax. One is 140.8 kb upstream and the other 13 kb downstream of the *Xbra*/*tbxtb* TSS (Extended Data Fig. 6 and 7); the transcriptional activities of these two CRMs have not yet been tested. Nevertheless, our results demonstrate that the SFZE syntax is present in both *brachyury* paralogs of vertebrates, reinforcing the idea of its ancestral association with *brachyury* orthologs and retention after 2R

## Discussion

Functionally equivalent enhancers have been discovered due to their sequence similarity^46,47^. Yet, enhancer sequence similarities tend to be significantly lower when comparing organisms across remote phylogenetic distances^47^. Previous studies have shown that enhancers without detectable sequence similarity could still be functionally equivalent^48,49^ ^50^. In this study, we show that the regulatory features of functionally equivalent enhancers can be preserved by specific motif grammar. We uncover a conserved SFZE syntax associated with *brachyury* of species across a wide range of taxa, including zebrafish, ascidian, hemichordate, sea urchin, fruit fly, sea anemone, and even a non-animal unicellular eukaryote. The syntax comprises binding sites for Su(H), Foxh1, Zic, and Ets, with the latter three arranged in close proximity and in a defined orientation and order. Our discovery of the *brachyury* SFZE in diverse organisms suggests that the deeply conserved regulatory code is likely under strong evolutionary constraints to maintain the specific syntax, as the adjacent sequences show extensive diversity.

At the onset of gastrulation, ambulacrarian and chordate *brachyury* genes are expressed preferentially on the ventral and dorsal side of the future blastopore, respectively. These positions can be considered comparable in light of the dorsoventral inversion that occurred in the chordate ancestor^51–53^. Our results showed that the SFZE-containing CRMs possess transcriptional activity in the zebrafish dorsal margin (part of the future blastopore) and in cells surrounding the sea urchin blastopore, suggesting that the SFZE syntax has a conserved function in the blastoporal region in deuterostomes. During gastrulation, chordate *brachyury* genes continue to be expressed in the dorsal axial region of the invaginating archenteron (i.e., the presumptive notochord), whereas ambulacrarian *brachyury* genes are only expressed in a few ventral endodermal cells (Extended Data Fig. 8). In several vertebrate and ascidian species, the initiation and continuity of *brachyury* expression in the notochord is orchestrated by Nodal and FGF signals. In the basal chordate amphioxus, the notochord expression of *brachyury* is controlled by Nodal signaling^54^, but it is not strongly affected when FGF signaling is inhibited^55^. Nevertheless, within the potential notochord enhancers of the two *brachyury* paralogs of the amphioxus *Branchiostoma floridae* (*Bfbra1* and *Bfbra2*), we found an SFZE motif in *Bfbra1* and two pairs of Su(H) binding sites in *Bfbra2* (Extended Data Fig. 6). These findings suggest that the SFZE syntax serves as a regulatory connection linking Nodal and FGF signals to control the sustained expression of *brachyury* in the notochord, and this function appears to have arisen at least before the divergence of tunicates and vertebrates (Fig. 5). Unlike the synergistic effect of the Foxh1 and Ets sites in the zebrafish notochord, the endodermal activity of the two TF binding sites during sea urchin gastrulation is temporally decoupled. Intriguingly, *foxh* orthologs could not be identified in the sea urchin and hemichordate genomes, suggesting that the gene was lost in the ambulacrarian lineage^56,57^. Additionally, inhibition of Nodal or FGF signals does not affect endodermal expression of sea urchin and hemichordate *brachyury*^58,59^ ^60^. Similarly, in *Drosophila*, *nodal* and *foxh1* orthologs have not been found in the genome^61–63^. It is likely that the Foxh1 sites in the SFZE motifs of ambulacrarian and *Drosophila* are occupied by other Fox TFs, which target similar core consensus sequences^64,65^. Therefore, the deeply conserved SFZE syntax embedded within the *brachyury* CRM may respond to different upstream factors in various lineages. We thus propose that the notochord could have evolved from rudimentary endodermal cells, and during chordate evolution, co-option of Nodal and FGF signals reinforced *brachyury* expression in the dorsal axial region of the invaginating archenteron. Further co-option of essential notochord differentiation genes downstream of *brachyury*^66^ would have also been necessary for evolution of the notochord.

**Figure 5.**
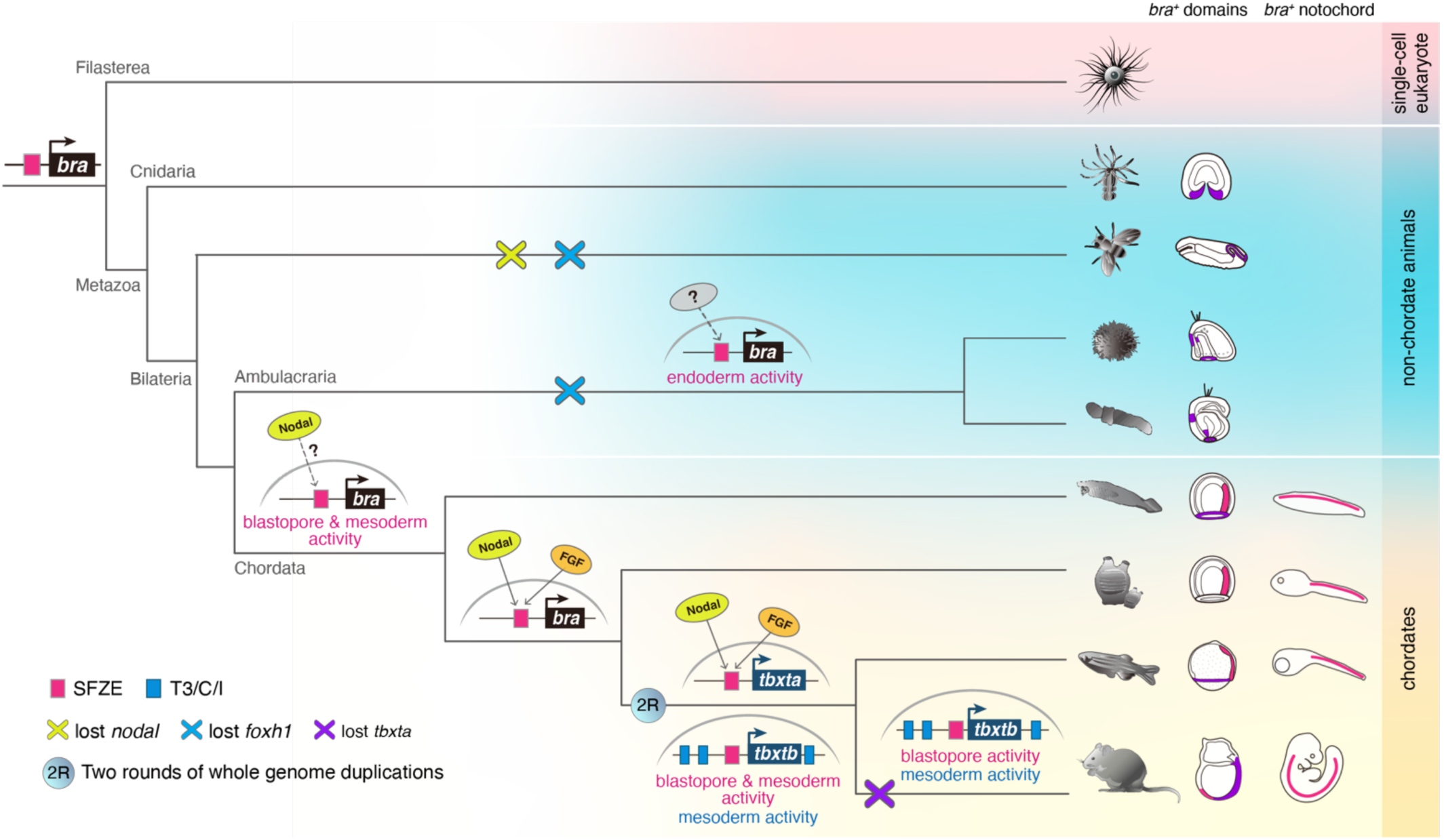
Proposed scenario of SFZE evolution and the origin of the notochord. The SFZE syntax (red rectangles) associated with *brachyury* orthologs could have already been present before the emergence of animals. The regulatory network that activates *brachyury* in the axial mesoderm by concurrent Nodal and FGF signals through the SFZE syntax may have been established in the chordate lineage, at least before the divergence of tunicates and vertebrates. The Ets and Fox family factors (activated by yet to be determined signaling pathways, indicated by question marks) likely regulate *brachyury* expression through SFZE in at least ambulacrarians. Some components of the regulatory inputs could have been modified due to the loss of *nodal* and/or *foxh* genes in *Drosophila* and ambulacrarians (cross symbols). After two rounds of whole genome duplications (2R), the SFZE syntax remained associated with both *brachyury* paralogs. Other notochord enhancers (T3/C/I) were subsequently evolved to control one of the paralogs (i.e., *tbxb*). In tetrapods, only *tbxb* was retained, and the notochord activity of the mammalian SFZE may have been substituted by T3/C/I enhancers.

Previous studies have revealed that the 5’ promoter regions of *Cibra* and *Drntl*/*tbxta* are sufficient for *brachyury* expression in the notochord. However, the corresponding regions of *tbxtb* in mouse^44^ and *Xenopus*^33,67^ lack notochord activity, instead driving expression in the primitive streak/blastopore. Several notochord-specific enhancers (T3, C and I) have been identified in the *tbxtb* loci of mouse^68^ and various jaw vertebrates^69^. These CRMs show sequence conservation across jaw vertebrates, and contain functional Brachyury binding sites for auto-regulation. However, T3, C and I enhancers are found neither in *tbxta* of ray-finned fishes nor the genomes of jawless vertebrates and *Ciona*, suggesting an origin in the last common ancestor of jawed vertebrates^69^. Notably, the notochord activity of T3, C and I is observed during early somitogenesis in zebrafish, after the initial expression of *Drbra*/*tbxtb* in the dorsal margin prior to gastrulation^43^. These findings suggest that the early expression of *Drbra*/*tbxtb* is regulated by another distinct enhancer, likely the SFZE-containing CRMs that we identified. Intriguingly, the proximal enhancer of the mouse *Mmbra* contains the SFZE syntax, but it exhibits activity in the primitive streak, and not in the notochord. Thus, although SFZE-containing CRMs are active in both the blastoporal region and notochord during zebrafish gastrulation, their notochord activity may have been fully substituted by T3, C and I enhancers in mammals.

Altogether, our study highlights the importance of conserved regulatory syntax in evolution of the notochord. The identified SFZE syntax has very deep evolutionary roots, likely originated before the emergence of animals. While the functions of the SFZE in *Capsaspora* remain to be determined, we propose that the ancestral SFZE syntax functions in endodermal cells during gastrulation. Our findings further suggest subsequent co-option of upstream factors and functional divergence of the SFZE syntax after duplication, which shaped its transcriptional activity during chordate evolution.

## Materials and Methods

### Animal collection and embryo culture

Adult sea urchin *S. purpuratus* were obtained from Amro Hamdoun (University of California, San Diego) and kept at 15°C. Mature sea star *A. typicus* and hemichordate *P. flava* were collected during the breeding season (June to August and September to December, respectively) from Chito Bay, Penghu Islands, Taiwan. Spawning and embryo cultures were conducted as previously described^70–72^ with modifications for *A. typicus*. In brief, starfish were weighed and injected with 1 mL of 200 *μ*M 1-methyladenine per 100 g weight on the dorsal side of arms to induce spawning. Starfish embryos were cultured at 33°C.

### Annotation of *brachyury* CRMs, promoters of *Pfbra* and *Spbra*, and the SFZE syntax

JBrowse^73^ (version 1.16.10) was used to visualize the presumed CRMs at the *brachyury* loci of zebrafish, *P. flava*, *N. vectensis*, and *C. owczarzaki*. The processed datasets of ATAC-seq of zebrafish and the histone ChIP-seq of *N. vectensis* and *C. owczarzaki* were obtained from NCBI Gene Expression Omnibus (GEO) under respective accession numbers GSE130944^31^, GSE46488^36^, and GSE71131^37^. The ATAC-seq dataset of hemichordate *P. flava* was obtained from a previous study and analyzed using standard pipelines^26^. The sequencing reads were aligned to the cognate genome assemblies of zebrafish (Zv10)^74^, *N. vectensis* (Nemvec1)^75^, *C. owczarzaki* (C_owczarzaki_V2, accession number in the European Nucleotide Archive: GCA_000151315.2, https://www.ebi.ac.uk/ena/browser/view/GCA_000151315.2), and *P. flava* (ptychodera_flava version 1.0.114)^76^ using Bowtie2 (version 2.5.1)^77^. The ChIP-seq or the ATAC-seq datasets of mouse, *Xenopus*, *Ciona*, sea urchin, and *Drosophila* were from the NCBI Genome Data Viewer^78^, Xenbase (v10.1)^45^, ANISEED^79–81^, Echinobase^34^, and FlyBase^35^, respectively. The promoters of *Pfbra* and *Spbra* were deduced by identifying positions of the core promoter elements (TATA box, initiator, downstream TFIIB recognition element, and downstream promoter element)^82,83^ around the transcription start site, as well as the promoter-proximal elements (CCAAT and GC box) upstream of the core promoter elements^84^. The SFZE motifs were identified by scanning the DNA sequences of putative CRMs to locate the binding sites of Zic and Foxh1, using the vertebrate PFMs (position frequency matrices) from the JASPAR database^85^. The Ets and Su(H) binding sites were detected according to the consensus core sequences MGGAW^86,87^ and GTGTRGAR^30^, respectively. The sub-optimal Su(H) binding site (Su(H)-like) was recognized as NTGRGAR or GTGRGAN

### Generation of GFP knock-in BAC clones and EGFP reporter constructs

The *Spbra*:*gfp* BAC (BAC clone ID: Sp_117A03_L in the Echinobase)^34,88^ contains an insert of ∼131 kb of DNA, with *Spbra* located 24 kb from the end. The genomic DNA library of *P. flava* was constructed in the pBACe3.6 vector in Eric Davidson’s laboratory at California Institute of Technology. To identify BAC clones containing *Pfbra*, BAC arrayed library filters were screened with a DIG-labeled *Pfbra* probe (PCR DIG probe synthesis kit, Roche) following the hybridization protocol (DIG high prime DNA labeling and detection starter kit II, Roche). The DNA probe spans from 3’ of exon 6 to the flanking intron with a size of 173 bp. The identified *Pfbra* BAC clone was obtained from the Echinobase (clone position: plate #57, well J10)^34^. The insert size of the *Pfbra* BAC was determined to be ∼146 kb by digestion with NotI-HF (NEB) and separation via pulse-field gel electrophoresis. End sequencing and mapping to the *P. flava* genome revealed that *Pfbra* is located ∼23 kb from the terminal end of the insert. To create a GFP knock-in BAC construct (*Pfbra*:*gfp* BAC), the *gfp* coding sequence was inserted at the TSS of *Pfbra* using homologous recombination^89,90^. NucleoBond® Xtra BAC plasmid purification kit was used to extract the BACs. Putative CRMs and promoters of *Pfbra* and *Spbra* were PCR amplified from the BACs. CRMs of zebrafish, fruit fly, and *N. vectensis* were PCR amplified from the cognate genomic DNA. CoCRM4 was synthesized by BIOTOOLS Co., Ltd. The CRMs and promoters were constructed into the HLC (Hugo’s lamprey construct) reporter vector^27,91^ by either restriction enzyme-mediated method, Gibson assembly (NEB), or combined by fusion PCR before ligating to the reporter vector. To modify particular TF binding sites in the SFZE syntax, the Q5® Site-Directed Mutagenesis Kit (E0554, NEB) was used to mutate the Foxh1 sites or the two guanine/cytosine nucleotides in the consensus core sequence (M<u>GG</u>AW) of the Ets sites. However, for the PfCRM2w/oFoxh1 construct, the 5’ end of the PfCRM2 was truncated by 224 bp to eliminate the two Foxh1 binding sites. The primer sets utilized in this study are listed in Extended Data Table 8..

### Microinjection

For microinjection of *P. flava*, zygotes were first pipetted for 10 s in filtered seawater (FSW) containing 0.25 % N-acetyl-L-cysteine (NAC, SIGMA) and incubated for 2 min to dissolve the jelly coat. The zygotes were then washed three times with FSW to remove NAC before being aligned on a protamine sulfate-coated culture dish for injection. The injection mixture used for *P. flava* included 5 ng/μl of linearized *Pfbra*:*gfp* BAC, 0.75 μg/μl Dextran Alexa Fluor™ 555 (Invitrogen), and 0.12 M KCl. Microinjection of *S. purpuratus* and *D. rerio* was conducted according to published procedures^92,93^. For sea urchins, 9 pl of injection solution containing either 5 ng/μl of linearized *Spbra*:*gfp* BAC or 0.4 ng/μl of PCR amplicons of reporter constructs, 0.75 μg/μl Dextran Alexa Fluor™ 555, and 0.12 M KCl was injected into the zygotes. Additionally, 15.4 ng/μl of carrier DNA (HindIII digested *S. purpuratus* genomic DNA) was added to the solution to facilitate the incorporation of PCR amplicons. For zebrafish, 2 nl of the injection solution containing 200 pg of circular reporter constructs and 0.03 %/μl of phenol red solution (SIGMA) in Danieau’s water was injected into the zygotes. When introducing BACs into zebrafish, 250 pg of circular BAC plasmids in 4 nl of injection solution was used. Reporter gene expression was observed and imaged with either a Zeiss Axio Observer Z1 or a Nikon SMZ18 microscope.

### Statistical analyses

Results of each transgenesis experiment in hemichordates, sea urchins and zebrafish are listed in the Extended Data Table 1-3 and 5-6. Statistical differences between activities of different CRMs or BACs were analyzed by the two-tailed Welch’s t-test with 95% confidence intervals. Bonferroni correction was applied to control Type I error rate for multiple comparisons, and *p-*values are shown in Figure 3-4. Statistical graphs were generated using GraphPad Prism 10.

### Whole-mount *in situ* hybridization

Antisense RNA probes were synthesized following the instructions of DIG RNA Labeling mix (Roche) or LabelIT DNP Labeling kit (Mirus) to generate respective DIG-labeled or DNP-labeled probes. *In situ* hybridization of *P. flava* and sea urchin embryos was conducted as described^53^. *A. typicus in situ* hybridization was performed according to the same protocol as used for sea urchins. Single and double FISH for sea urchin, *P. flava*, and zebrafish was performed following the previously described procedure^93–95^, except that 100 mM sodium azide was used to quench endogenous peroxidase activity or antibody activity. Tyramide Signal Amplification (TSA; PerkinElmer) was applied to amplify the fluorescent signals. Embryos were imaged with either a Zeiss Axio Imager A2 or a Zeiss Axio Observer Z1 microscope. Fluorescent signals were captured using a Zeiss LSM 880 confocal microscope.

## Acknowledgments

We thank the Taiwan Zebrafish Core Facility at Academia Sinica (TZCAS) (NSTC 112-2740-B-400-001) for providing the zebrafish ASAB wild-type strain. We are grateful to Sheng-Ping L. Hwang, Hwei-Jan Hsu, and Hiroshi Watanabe for providing genomic DNA of zebrafish, fruit fly, and *N. vectensis*. We also thank Christopher Lowe for sharing the HCL reporter vector. We thank Yu-Fen Lu for technical assistance in zebrafish microinjection. We also thank Marcus Calkins for English editing. We are grateful for the help from the core facility and the Marine Research Station of the Institute of Cellular and Organismic Biology, Academia Sinica. This study was supported by National Science and Technology Council, Taiwan (NSTC-113-2326-B-001-004 and NSTC-113-2811-B001-093).

## Author contributions

TPF and YHS conceived the project. TPF designed and performed the experiments. TPF and YHS wrote the manuscript. CYL processed the ATAC-seq and ChIP-seq data. JRL and YCC were involved in *in-situ* hybridization experiments. JRL was involved in microinjection of *Spbra* BAC into sea urchins. RAC and AEC initiated CRM analysis of the sea urchin *brachyury* gene. YHS and JKY supervised the project. All authors read and approved the final manuscript.

## Additional information

Correspondence and requests for materials should be addressed to YHS.

---

**Extended Data Figure 1.**
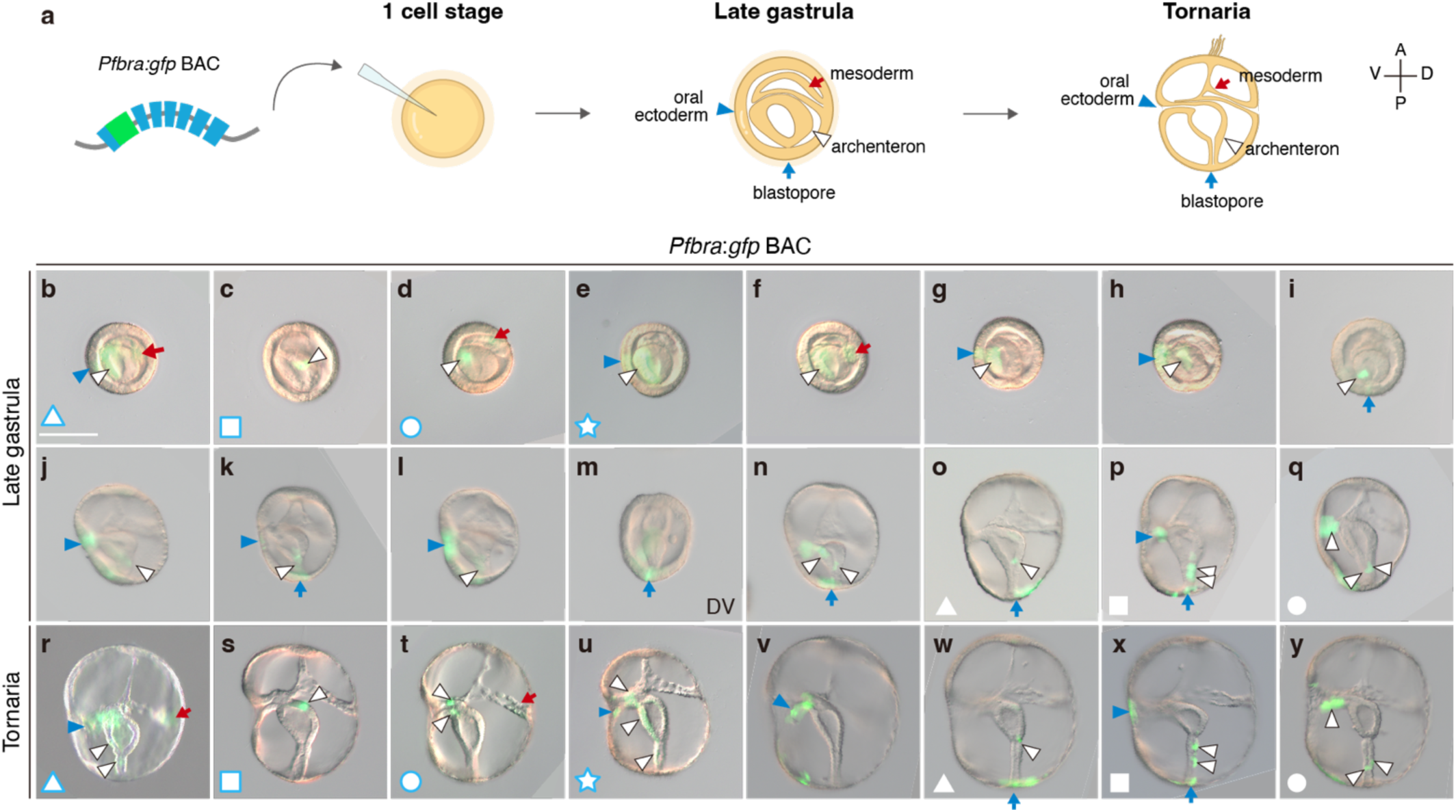
*Pfbra:gfp* BAC recapitulates the endogenous expression pattern of *Pfbra*. **a**, *P. flava* zygotes injected with linearized *Pfbra*:*gfp* BAC were observed at the late gastrula and tornaria stages (A, anterior; P, posterior; V, ventral; D, dorsal). **b-y**, Images of embryos from 5 independent experiments (Extended Data Table 1). The same embryos observed at different developmental stages are marked with the same symbols in the bottom left corners. On average, 30.1% of the injected embryos showed GFP signals, which were observed mainly in the oral ectoderm (blue arrowhead), archenteron (white arrowhead), and blastopore (blue arrow) at both late gastrula (b-q, 38-48 hpf) and tornaria (r-y, 72 hpf) stages. Ectopic expression in the mesoderm (17.6%) is indicated by red arrows. Embryos are viewed from the lateral side with the mouth to the left, unless otherwise indicated (DV, dorsal view, in panel m). All panels are shown at the same scale, according to the scale bar (100 μm) in panel b.

**Extended Data Figure 2.**
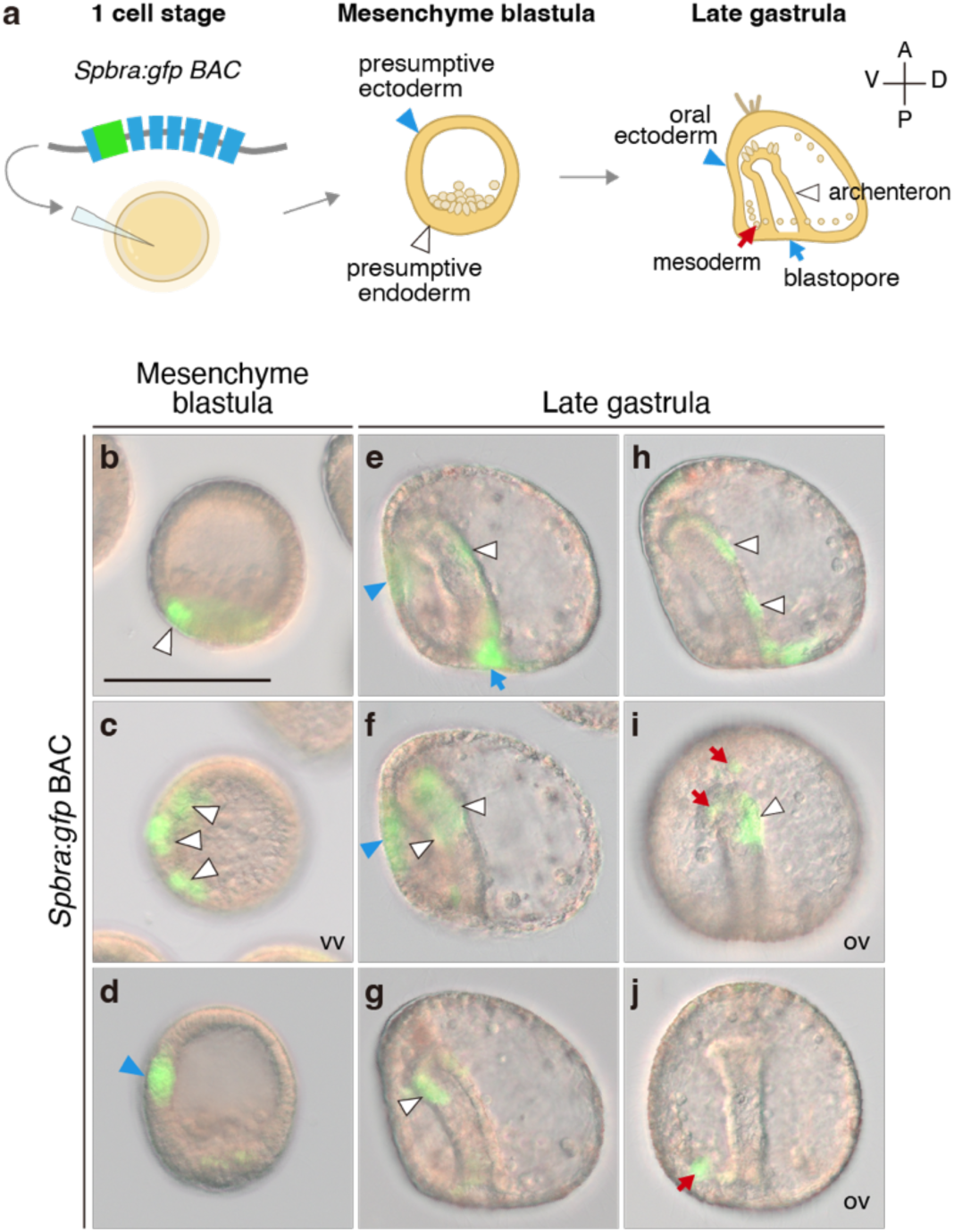
*Spbra:gfp* BAC recapitulated the endogenous expression pattern of *Spbra*. **a**, *S. purpuratus* zygotes injected with linearized *Spbra*:*gfp* BAC were observed at the mesenchyme blastula and the late gastrula stages (A, anterior; P, posterior; V, ventral; D, dorsal) (Extended Data Table 1). **b-d,** GFP signals were observed in the presumptive endoderm (white arrowhead) and ectoderm (blue arrowhead) at the mesenchyme blastula stage. Panel c shows the vegetal view of the embryo in panel b, with GFP signals present on one side of the embryo. **e-j,** At the late gastrula stage, GFP signals were observed in the oral ectoderm (blue arrowhead), blastopore (blue arrow), and archenteron (white arrowhead). Ectopic expression was also observed in the mesodermal cells (red arrow). Unless otherwise indicated, embryos are viewed from the lateral side (VV, vegetal view; OV, oral view). All panels are shown at the same scale, according to the scale bar (100 μm) in panel b.

**Extended Data Figure 3.**
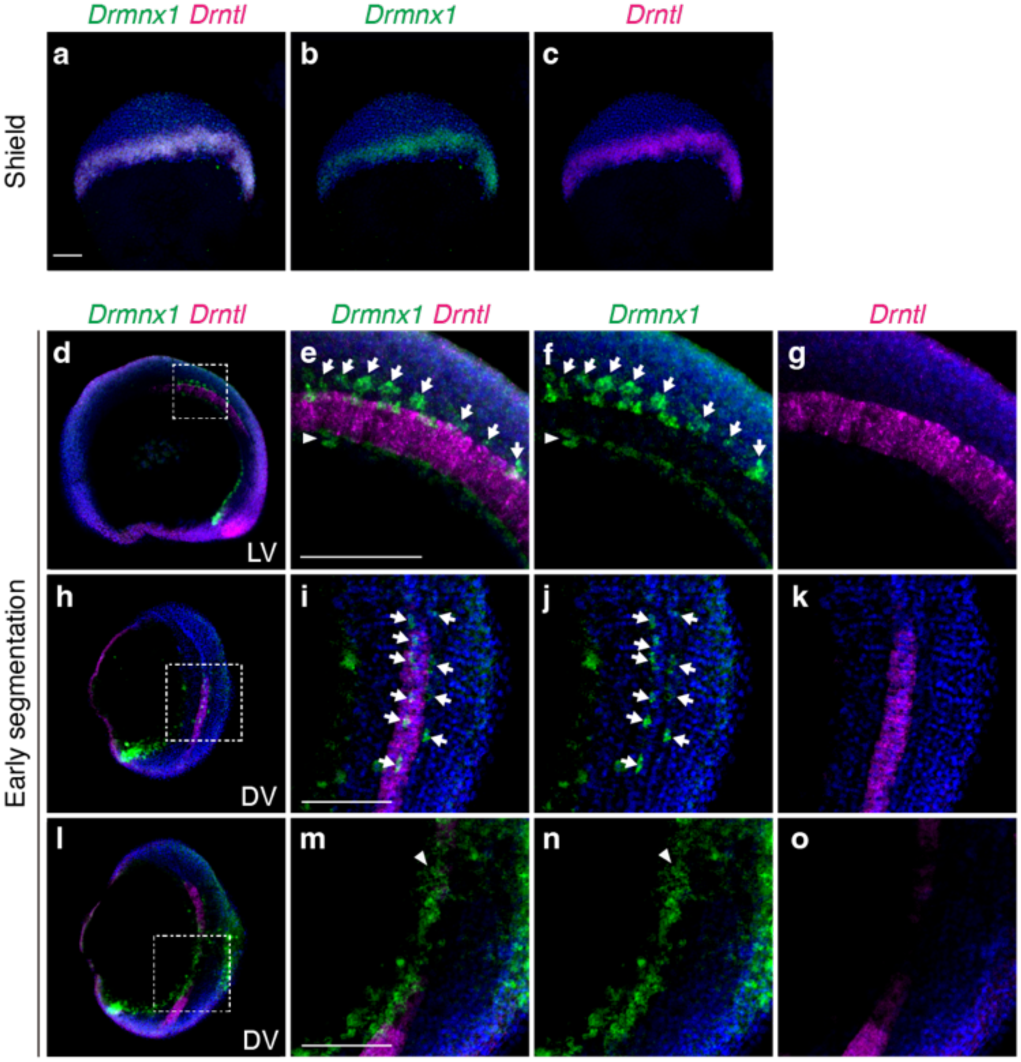
Expression of *Drmnx1* in motor neurons and hypochord of zebrafish embryos. Double FISH of *Drmnx1* (green) and *Drntl* (magenta) in zebrafish embryos. Nuclei were counterstained with Hoechst 33342 (blue). **a-c,** *Drmnx1 and Drntl* transcripts were detected in the germ ring at the shield stage. **d-o,** *Drmnx1* expression in the motor neurons and hypochord at the early segmentation stage. Panel d is a lateral view with dorsal side to the right. Panels h and l show the same embryo (dorsal side to the right), with different Z-stacks focusing respectively above and underneath the notochord. The white dashed boxes in panels d, h, and l are magnified and split into single channels shown in panels e-g, i-k, and m-o, respectively. *Drmnx1* expression in the motor neurons (white arrow) and hypochord (white arrowhead) is indicated. All scale bars represent 100 μm. Panels a-d, h, and l are in the same scale, and panels e-g, i-k, and m-o are in the same scale.

**Extended Data Figure 4.**
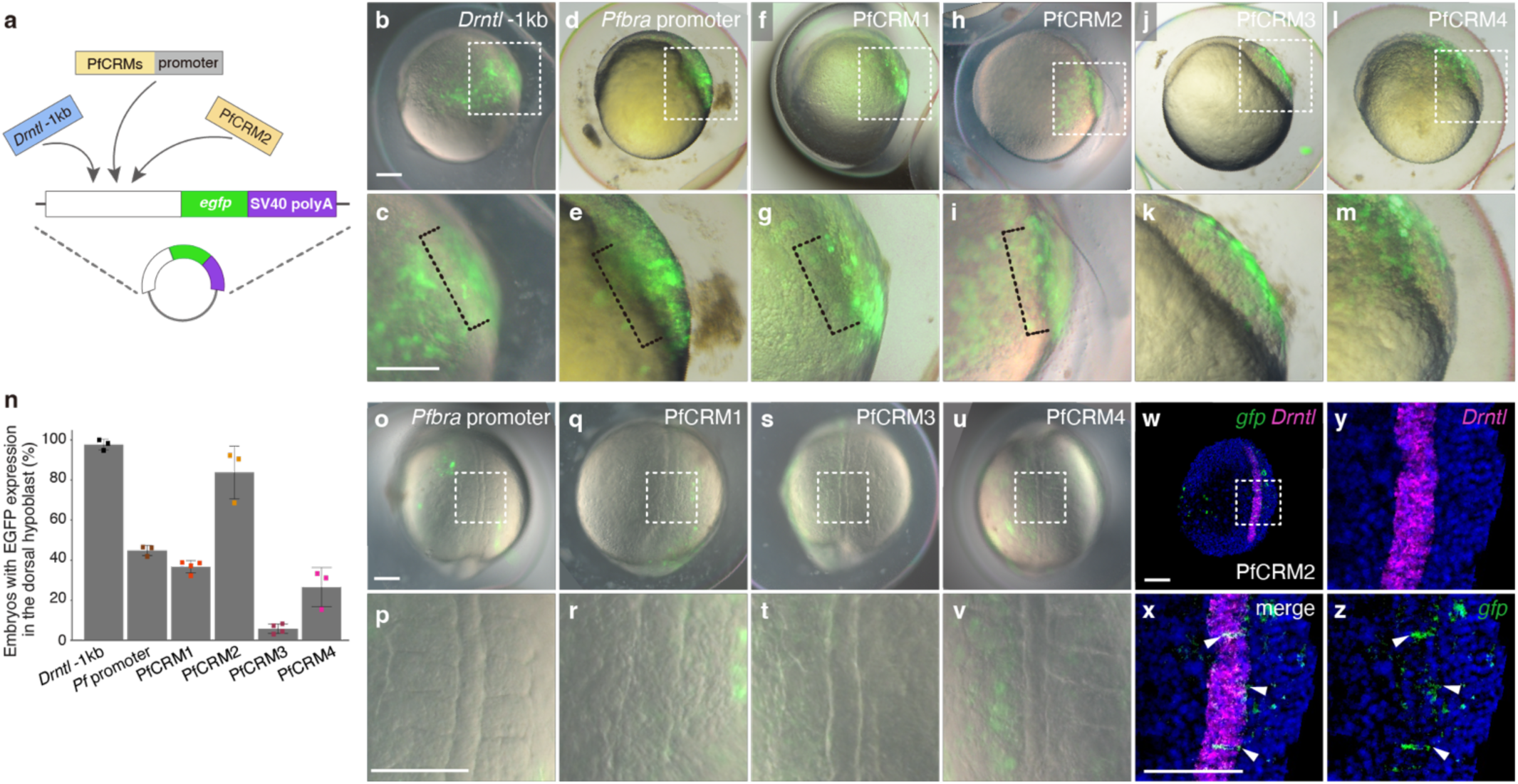
Transcriptional activities of zebrafish and hemichordate reporter constructs in zebrafish embryos. **a**, The illustration depicts structures of the reporter constructs. **b-m**, Activities of the *Drntl* notochord enhancer, *Pfbra* promoter, and PfCRMs in zebrafish embryos at the shield stage. Embryos are displayed in lateral view with dorsal to the right. Magnified views of the shield regions (white squares) are displayed in the lower panels. Black brackets highlight EGFP signals in the dorsal hypoblast (a layer of involuting cells that primarily give rise to the notochord). Embryos in panels k and m exhibit EGFP signals in the dorsal epiblast (cells that contribute to both notochord and the floor plate), but not in the hypoblast. **n**, The chart displays percentages of embryos exhibiting EGFP signals in the dorsal hypoblast out of the total number of EGFP-positive embryos. Each data point (colored square) is the result of a single experiment. The gray columns are average results from at least three biological replicates, with error bars showing standard deviations. **o-v**, Activities of *Pfbra* promoter and PfCRMs in zebrafish embryos at the early segmentation stage. Embryos are shown in the dorsal view. The notochord regions of each embryo are enlarged and displayed below the respective image. **w-z**, Double FISH of *Drntl* (magenta) and *egfp* (green) in zebrafish embryos injected with the PfCRM2 reporter. Nuclei were counterstained with Hoechst 33342 (blue). The early segmentation stage embryo is oriented in the dorsal view (slightly toward the right side). The notochord region in panel w (white square) is enlarged (x) and split into single channels (y-z). White arrowheads indicate *egfp* expression in notochord cells. Upper panels of b-m, lower panels of b-m, panels o, q, s and u, panels p, r, t and v, and panels x-z are in the same scale, respectively. All scale bars represent 100 μm.

**Extended Data Figure 5.**
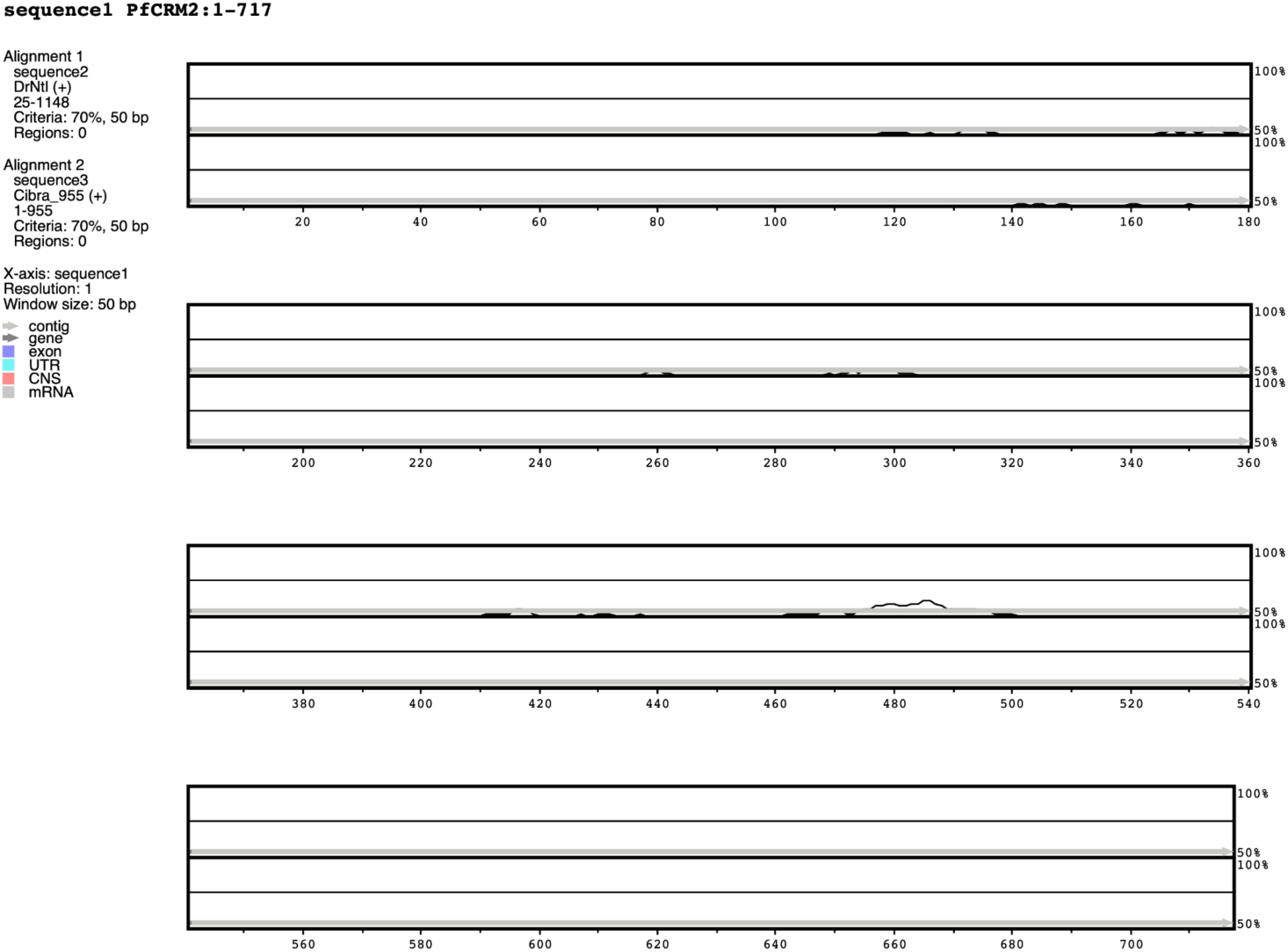
No conserved regions are detected between PfCRM2 and the notochord enhancers of zebrafish and *Ciona*. Sequence comparison using mVISTA with PfCRM2 as a query to align with the known notochord enhancers of zebrafish *Drntl* and *Ciona Cibra*. The criterion of 70% conservation in a 50 bp sliding window was used. Regions with 50-100% identity are displayed. The conserved regions should be marked with red peaks, but none are observed in the plot.

**Extended Data Figure 6.**
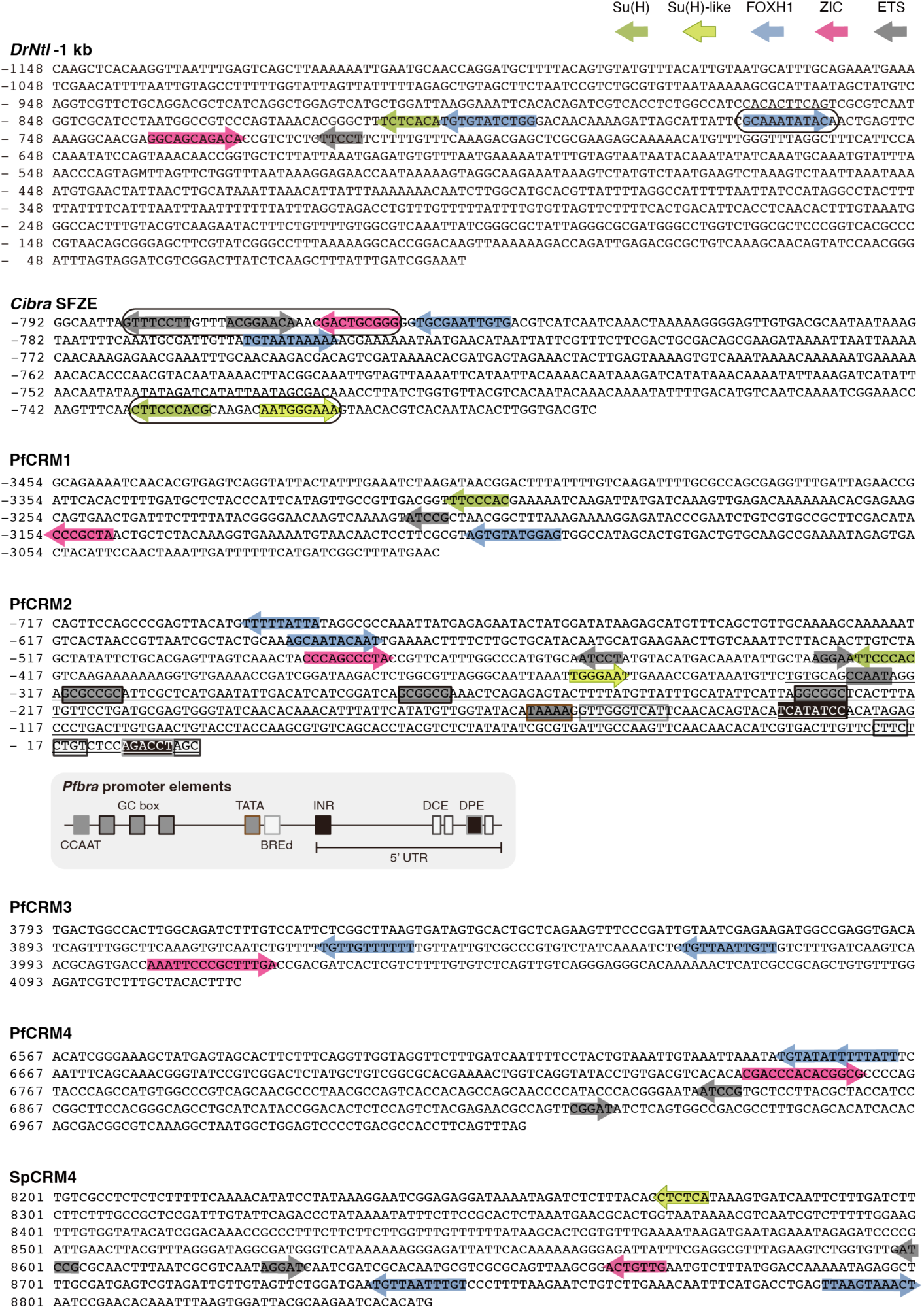
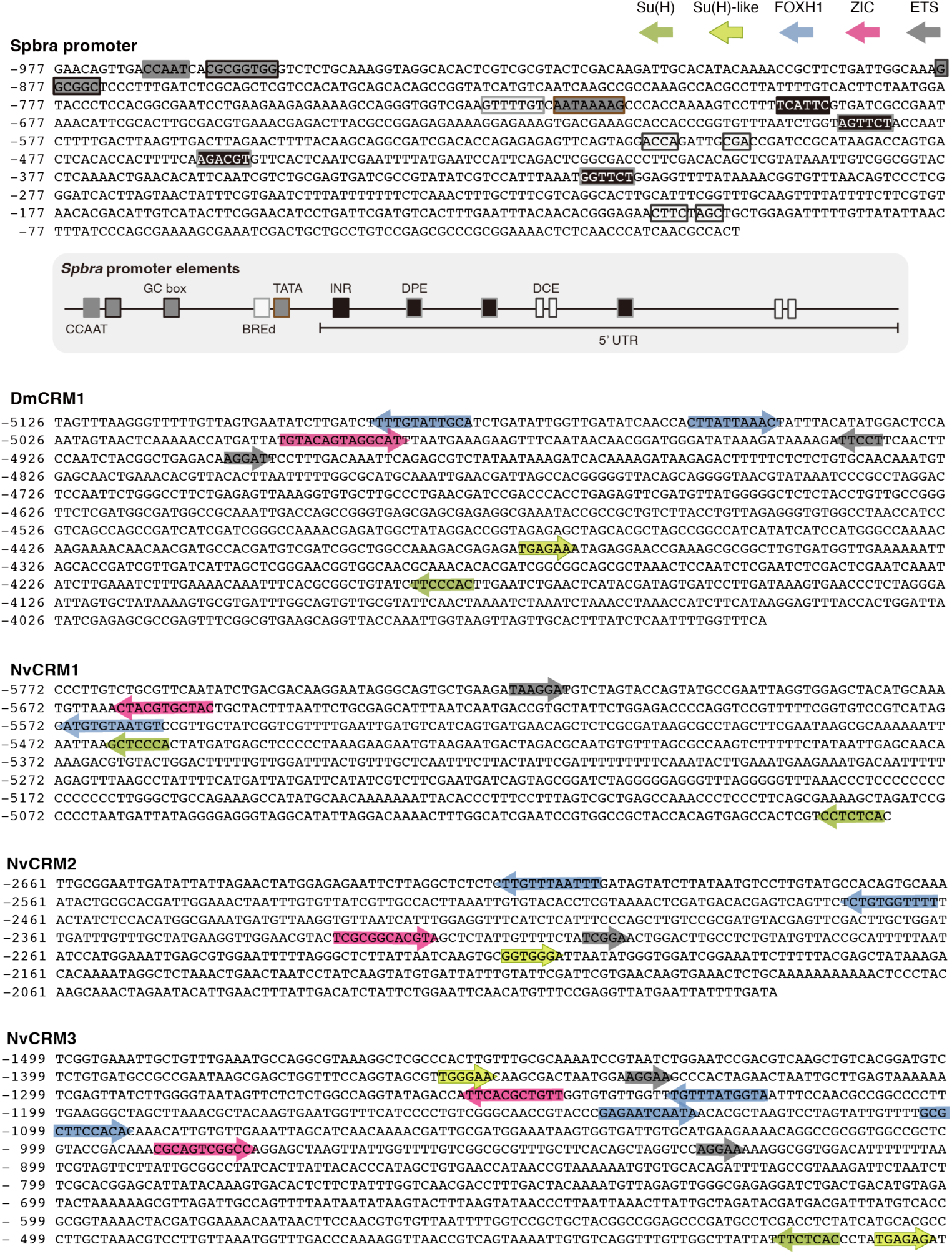
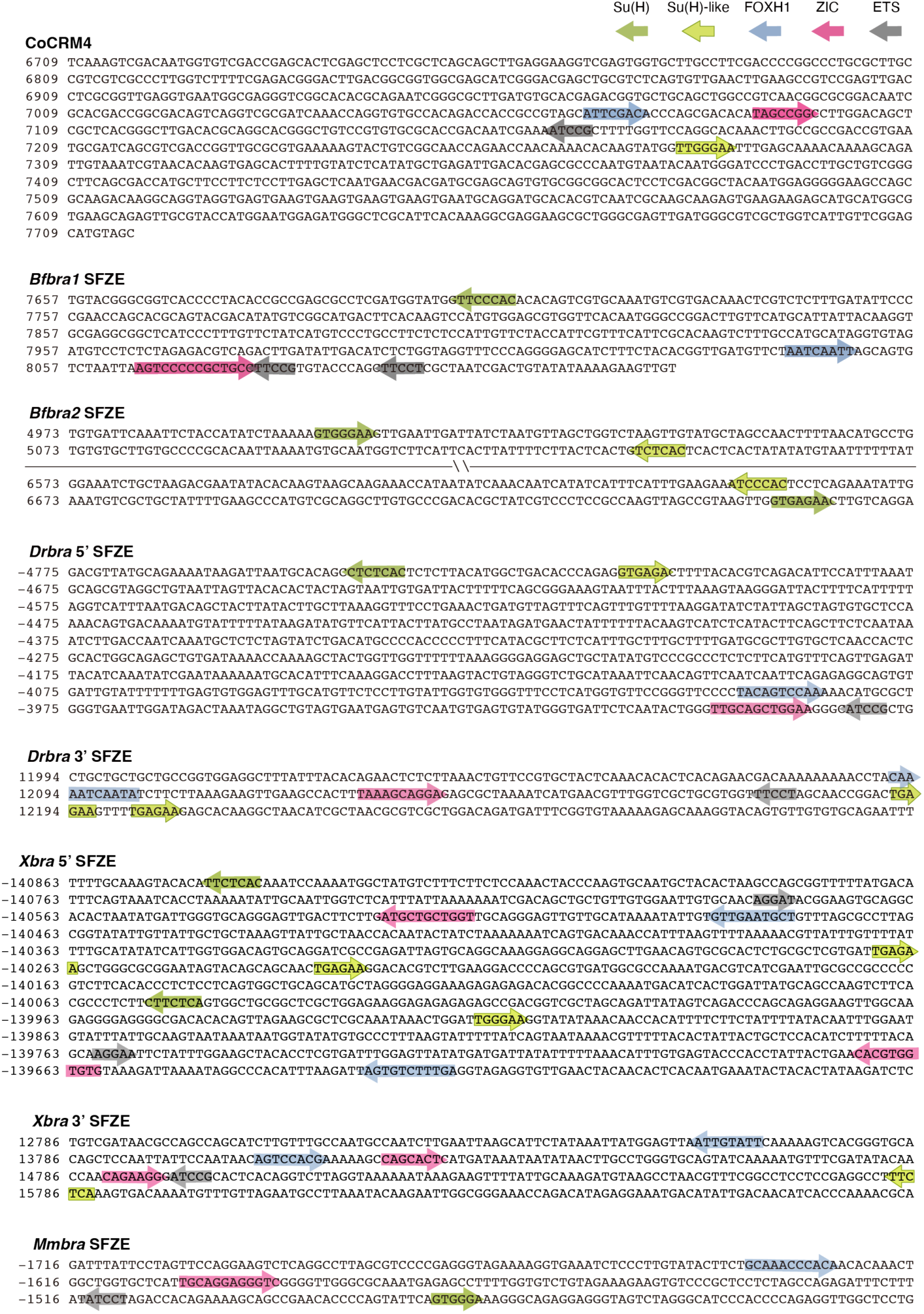
Sequences of *Pfbra*/*Spbra* promoters and SFZE/SFZE-like-containing CRMs/regions from diverse organisms. Positions of CRMs are labeled on the left side of the sequences, either upstream or downstream of the TSS of the examined *brachyury* orthologs. Binding sites of the four TFs are marked by colored arrows. Previously identified TF binding sites in the notochord enhancers of *DrNtl* and *Cibra* are circled. The sequence of the *Pfbra* putative promoter within PfCRM2 is underlined. The promoter/promoter-proximal elements within PfCRM2 and of *Spbra* promoter are enclosed by rectangles. The illustrations underneath PfCRM2 and *Spbra* promoter sequences depict the arrangements of the deduced promoter elements. BERd: downstream TFIIB recognition element; INR: initiator; DCE: downstream core element; DPE: downstream promoter element.

**Extended Data Figure 7.**
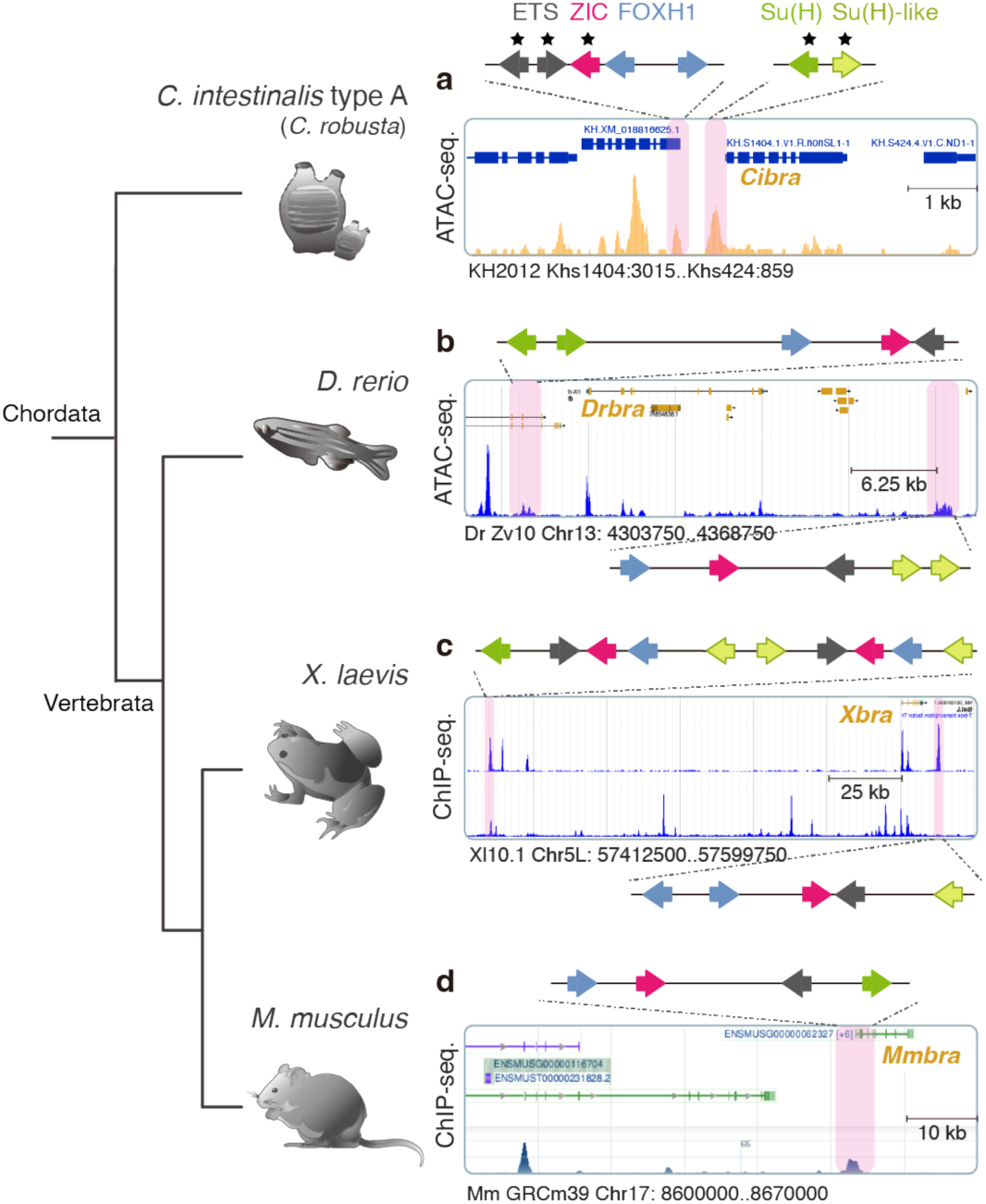
SFZE-containing CRMs in the *brachyury* loci of chordates. **a-b,** The open chromatin regions were revealed by ATAC-seq at the *Cibra* and *Drbra* loci. Previously identified TF binding sites in the notochord enhancer of *Cibra* are indicated by the black asterisks. **c-d**, Tracks display ChIP-seq of histone methylation (H3K4me3: upper track of c and the track in d) and Ets (lower track in c) at the *brachyury* loci of frog and mouse. CRMs highlighted by pink rectangles harboring the SFZE syntax. Orientations and orders of the TF binding sites are illustrated with colored arrows on top of the respective diagrams. The color key is indicated in panel a. Genome version, scaffold/chromosome number, and position of the genomic loci are displayed below the respective panel. Developmental stages or tissues/cells used for the ATAC-seq or ChIP-seq analyses are listed in Extended Data Table 7. Illustrations of *Xenopus* and mouse are adapted from BioRender.com.

**Extended Data Figure 8.**
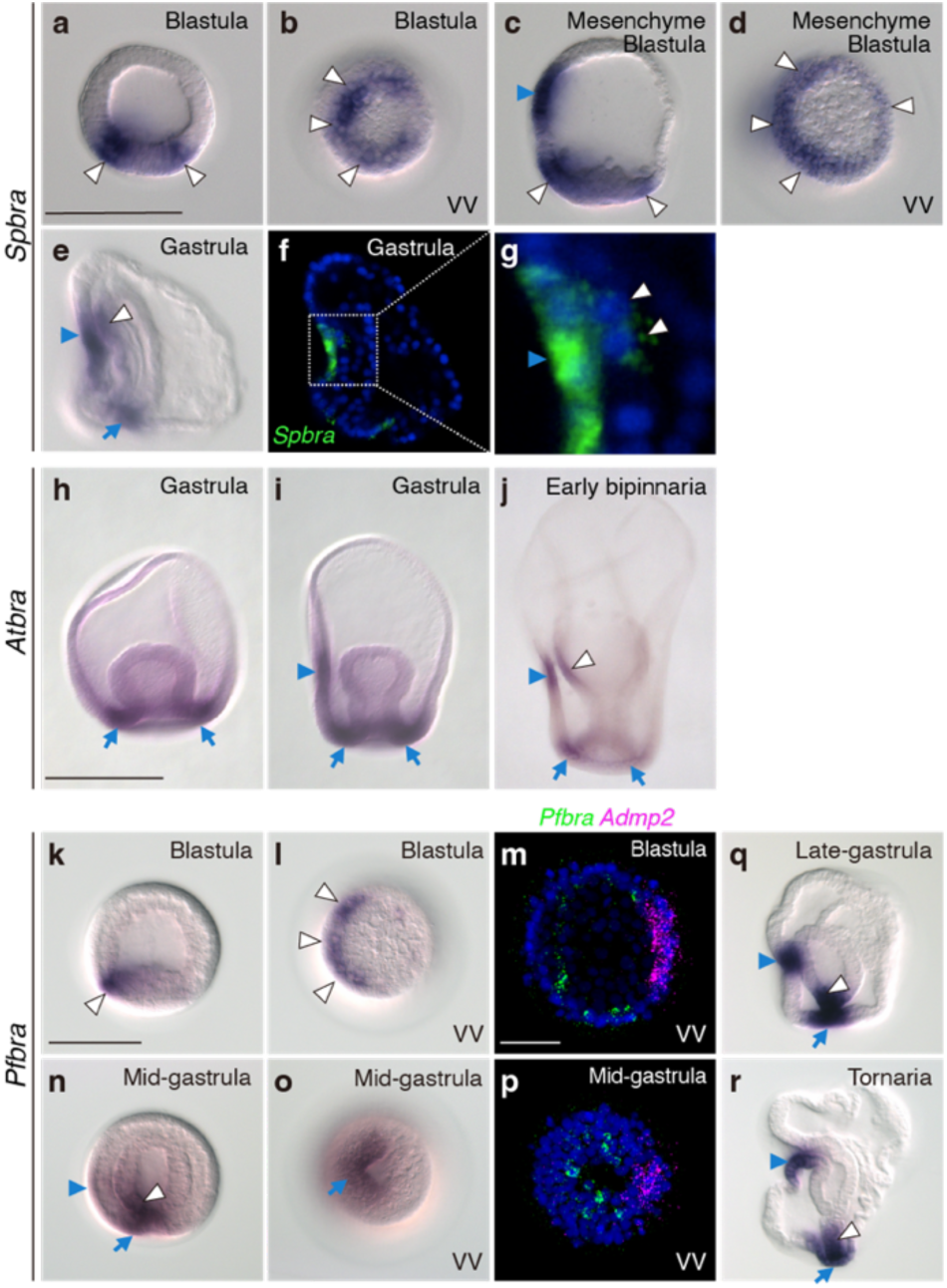
Expression of *brachyury* orthologs in ambulacrarians. **a-g**, As previously described^19^, during sea urchin embryogenesis, *Spbra* expression was initially detected in the vegetal region (white arrowhead) of the blastula (a-b), and an additional expression domain was observed in the oral ectoderm (blue arrowhead) at the mesenchyme blastula stage (c-d). Panels b and d display the vegetal view of the same embryos in panels a and c, respectively, revealing enhanced *Spbra* expression on one side of the vegetal plate. The side with enhanced *Spbra* expression is the oral (ventral) side, judging from its known oral ectodermal expression. **e**, *Spbra* expression was present in the oral ectoderm, cells surrounding the blastopore (blue arrow), and a few cells in the ventral archenteron (white arrowhead) at the late gastrula stage. **f-g**, FISH of *Spbra* (green). The oral region of panel f is magnified in panel g, highlighting *Spbra* expression in the ventral archenteron (white arrowhead). **h-j,** Expression of *Atbra* in the sea star *Archaster typicus*. The expression domain in the cells near the blastopore (blue arrow) and the oral ectoderm (blue arrowhead) during gastrulation. An additional expression domain in the ventral archenteron was detected at the early bipinnaria stage (white arrowhead). **k-r**, *Pfbra* expression during hemichordate embryogenesis. *Pfbra* transcripts are detected on one side of the vegetal region (white arrowhead) at the blastula stage (k-l). By the mid-gastrula stage (n-o), *Pfbra* expression appears in the oral ectoderm (blue arrowhead), with stronger expression observed in the ventral posterior archenteron (white arrowhead) and the ventral blastopore (blue arrow). Vegetal view of the panels k and n are displayed respectively in panels l and o. Double FISH of *Pfbra* (green) with a dorsal marker *Pfadmp2*^96^ (magenta) confirms that the asymmetrical vegetal expression is on the ventral side (m and p). As previously described^20^, *Pfbra* is expressed in the hindgut, oral ectoderm, and blastopore at the late gastrula and tornaria stages (q-r). Nuclei were counterstained with Hoechst 33342 (blue) in panels f, g, m, and p. All scale bars represent 100 μm. Panels a-g, panels h-j, panels k-l, n-o and q-r, and panels m-p are in the same scale, respectively. VV: vegetal view.

**Extended Data Table 1.** Reporter activities of BACs in hemichordate and sea urchin embryos.

| *P. flava* | GFP<sup>+</sup> |  | Injected |
| --- | --- | --- | --- |
|  | embryos (%) |  | embryos |
| *Pfbra:gfp* BAC | 5 | 25.0% | 20 |
|  | 2 | 11.8% | 17 |
|  | 4 | 57.1% | 7 |
|  | 5 | 45.5% | 11 |
|  | 1 | 11.1% | 9 |

| Mesenchyme blastula |  |  |  |  |  |  |  | Late gastrula |  |  |  |  |  |  |
| --- | --- | --- | --- | --- | --- | --- | --- | --- | --- | --- | --- | --- | --- | --- |
| *S. purpuratus* | Presumptive ectoderm (%) |  | Presumptive endoderm (%) |  | Mesenchyme cells (%) |  | GFP<sup>+</sup> embryos | Oral ectoderm & Blastopore (%) |  | Archenteron (%) |  | Mesenchyme cells (%) |  | GFP<sup>+</sup> embryos |
| *Spbra:gfp* BAC | 0 | 0.0% | 48 | 100.0% | 0 | 0.0% | 48 | 36 | 85.7% | 40 | 95.2% | 6 | 14.3% | 42 |
|  | 4 | 5.9% | 67 | 98.5% | 0 | 0.0% | 68 | 8 | 11.3% | 71 | 100.0% | 9 | 12.7% | 71 |
|  | 10 | 10.2% | 93 | 94.9% | 0 | 0.0% | 98 | 37 | 60.7% | 55 | 90.2% | 5 | 8.2% | 61 |

**Extended Data Table 2.**
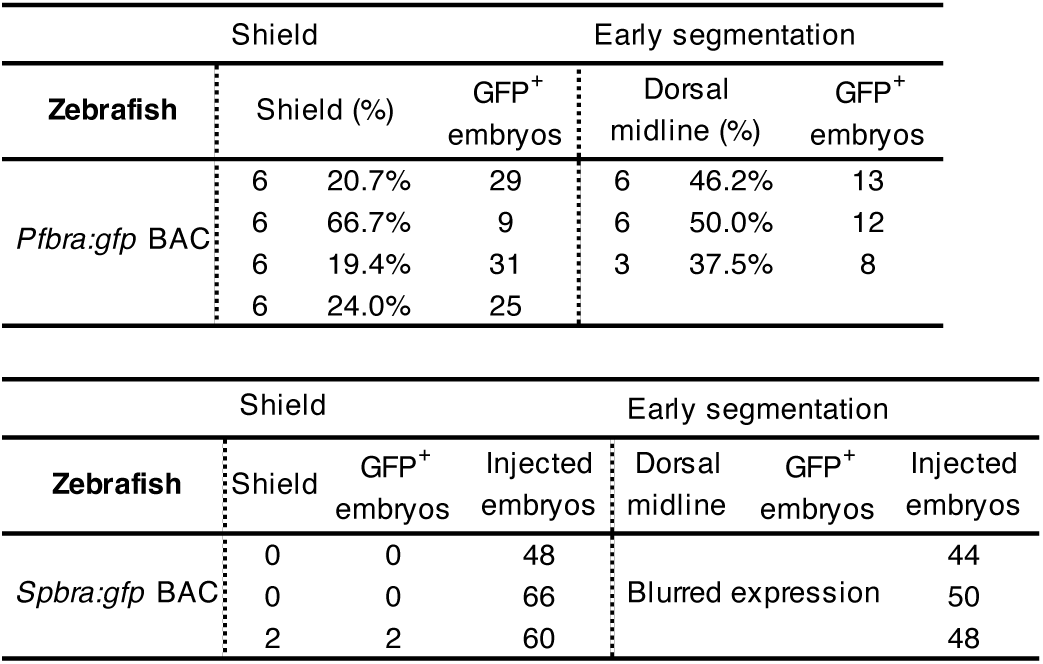
Reporter activities of BACs in zebrafish embryos.

**Extended Data Table 3.**
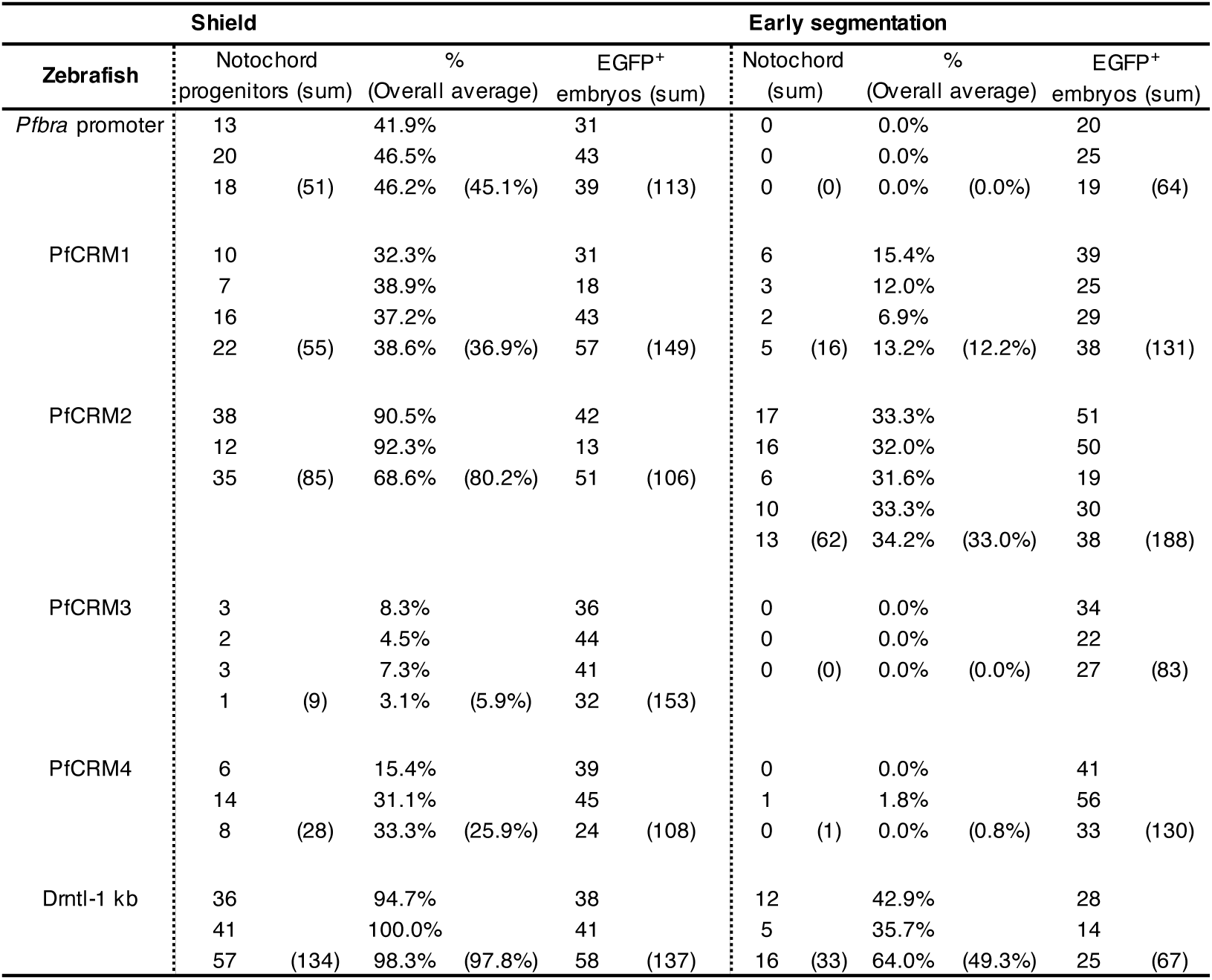
Reporter activities of PfCRMs in zebrafish embryos.

**Extended Data Table 4.**
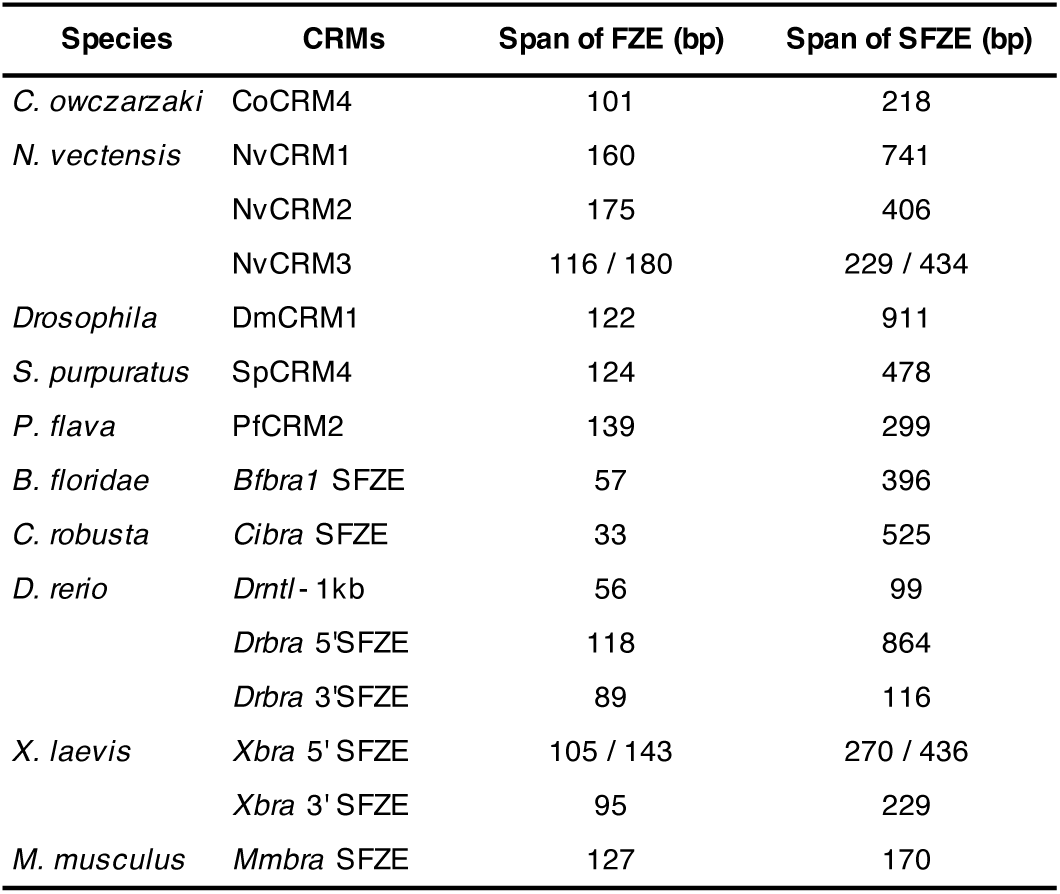
Spans of FZE and SFZE in various organisms.

**Extended Data Table 5.**
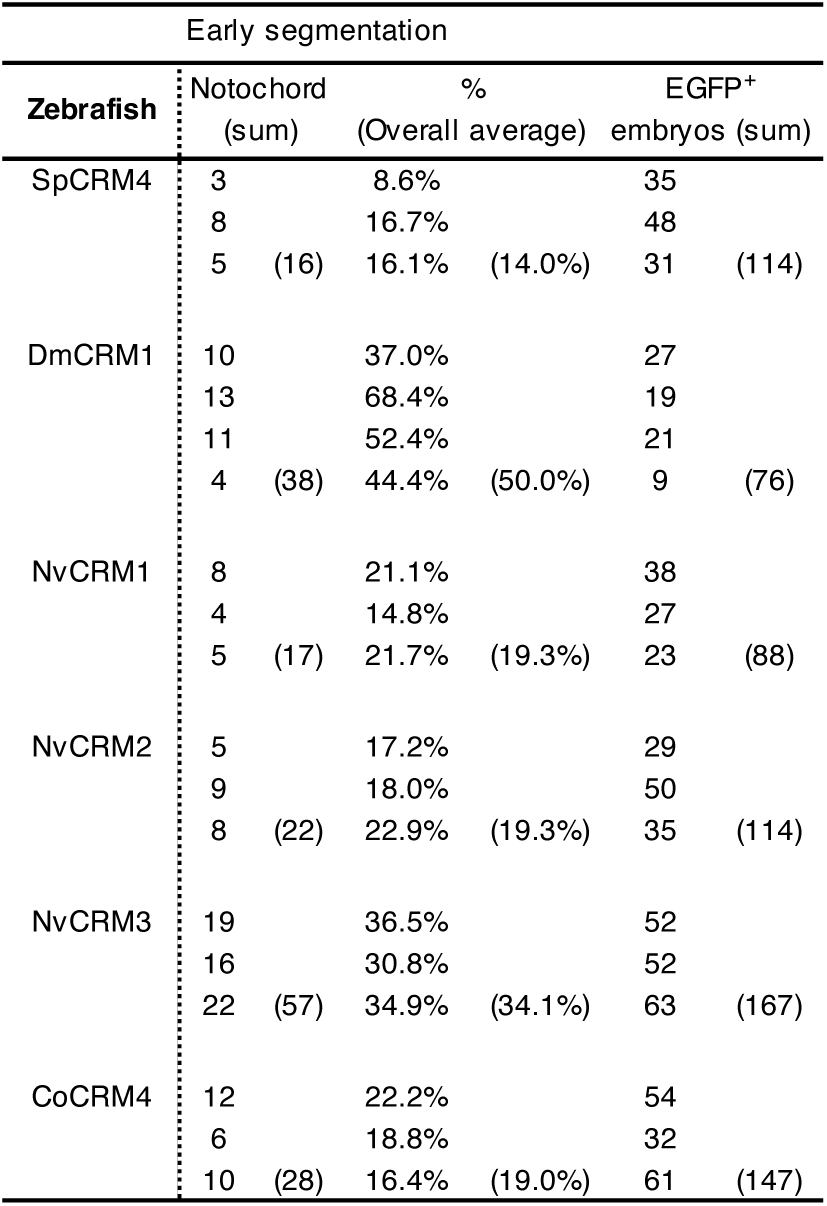
Notochord activities of SFZE-containing CRMs in zebrafish embryos.

**Extended Data Table 6.**
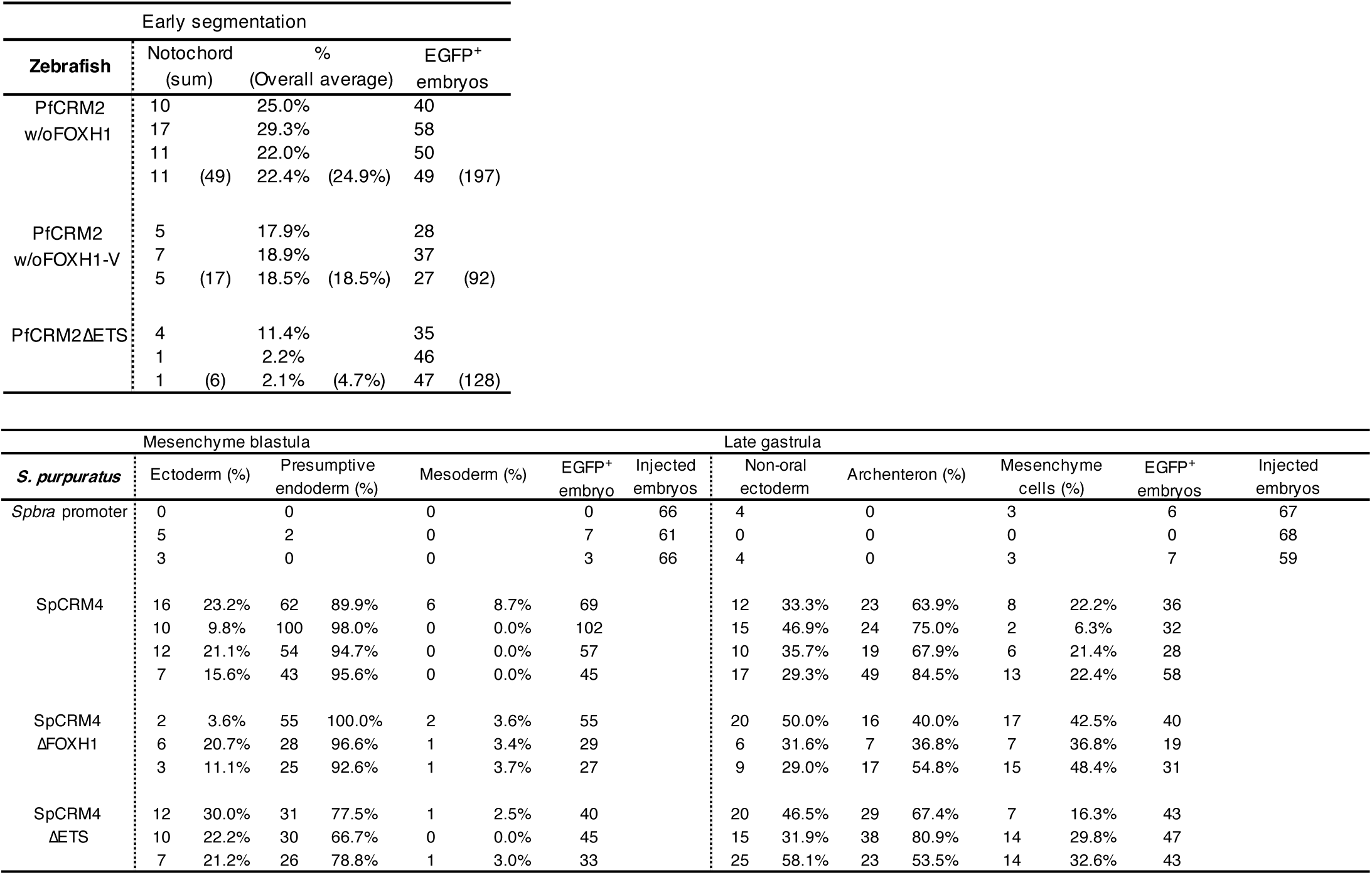
Activities of reporters without Foxh1 or Ets binding sites in zebrafish and sea urchin embryos.

**Extended Data Table 7.** ATAC-seq and ChIP-seq datasets used in this study.

| Organisms | Analytic type | Developmental stages/ Tissues/ Cells | Reference |
| --- | --- | --- | --- |
| *C. owczarzaki* | ChIP-seq | Filopodia | 44 |
| *C. robusta* | ATAC-seq | Late gastrula | 92-94 |
| *D. rerio* | ATAC-seq | 80% epiboly | 38 |
| *X. laevis* | ChIP-seq | NF 10.5 and NF 11.5 | 40 |
| *M. musculus* | ChIP-seq | Primary tissues, primary cells and immortalized cell lines | 91 |
| *P. flava* | ATAC-seq | Early blastula | 32 |
| *S. purpuratus* | ATAC-seq | Late gastrula | 41 |
| *Drosophila* | ChIP-seq | Embryo 6-8 h | 42 |
| *N. vectensis* | ChIP-seq | Gastrula | 43 |

**Extended Data Table 8.**
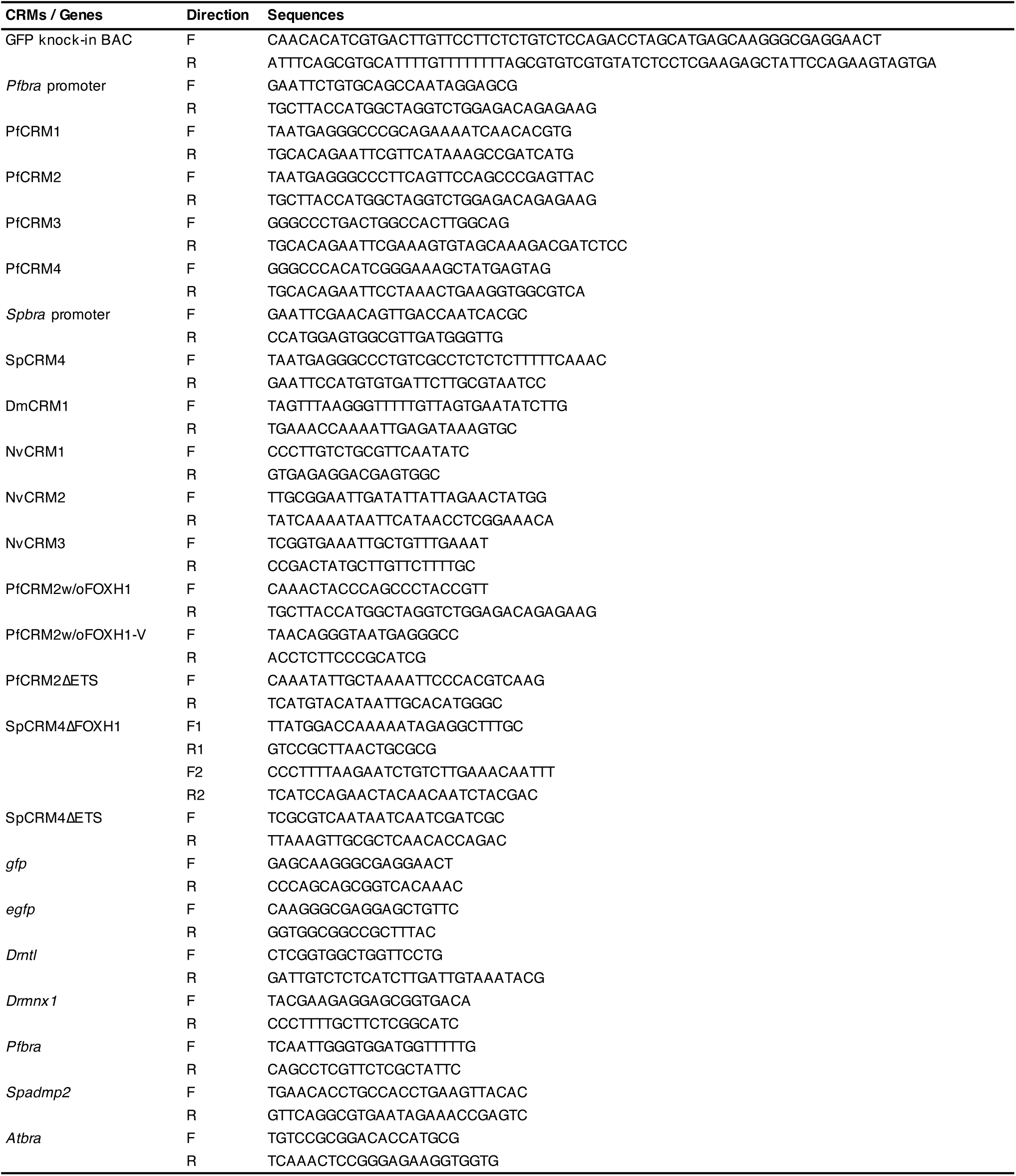
Primers used to generate reporter constructs and *in situ* hybridization probes.

